# Overcoming impaired antigen presentation in tumor draining lymph nodes facilitates immunotherapy

**DOI:** 10.1101/2025.09.17.676884

**Authors:** Meghan J. O’Melia, Lutz Menzel, Pin-Ji Lei, Hengbo Zhou, Neian Contreras-Alvarado, Johanna J. Rajotte, Lingshan Liu, Mohammad R. Nikmaneshi, James W. Baish, Jessalyn M. Ubellacker, Genevieve M. Boland, Sonia Cohen, Lance L. Munn, Timothy P. Padera

## Abstract

Immunotherapies have revolutionized cancer care in recent decades, but approved therapies often fail and currently only target specific steps in the generation of anti-cancer immune responses. Notably, the majority of approved immunotherapies do not target antigen processing and presentation, which are key steps in the development of immune responses and harbor potential as targets to improve immunotherapy. Here, we demonstrate that breast tumors induce locoregional lymph node impairment in antigen presentation—but not in antigen processing—which limits anti-cancer antigen- specific T cell responses. Inhibition of the locoregional T cell response was due to a tumor-mediated reduction of the cytokine IL1β in tumor draining lymph nodes, which impaired antigen presentation. Further, we tested the ability of dendritic cells in lymph nodes at various distances from the primary tumor to be activated utilizing an antigen- agnostic adjuvant delivery strategy. We observed improved anti-tumor T cell responses when the adjuvant was delivered to cancer antigen-positive lymph nodes distant from the tumor, suggesting these lymph nodes can be targeted to improve anti-cancer immune responses. When combined with immune checkpoint blockade, delivery of the adjuvant to distant lymph nodes led to long-term survival and protection from recurrence. Antigen presentation and T cell responses could also be recovered by exogenous delivery of IL1β via intratumoral injection, with improved survival when combined with immune checkpoint blockade. Our study demonstrates that tumor-induced locoregional impairment of antigen presentation can be overcome by the appropriate introduction of immunological adjuvant to tumor-distant lymph nodes or by restoring IL1β to the tumor-draining lymph node. These strategies can induce high-quality, durable immune responses and have clinical implications for expanding the efficacy of immunotherapies.

**Summary:** Breast tumors induce locoregional lymph node impairments in antigen presentation, which can be remedied via either IL1β delivery or antigen-agnostic adjuvant therapies to distant lymph nodes, facilitating better immunotherapy responses.

## INTRODUCTION

Breast cancer is the most commonly diagnosed malignant cancer in women. In the United States, 1 in 8 women will be diagnosed with breast cancer during their lifetime [1]. Fortunately, many breast cancers are detected early through screening programs, however treatments for late-stage breast cancer are currently insufficient, with only 31% of advanced breast cancer patients surviving long-term [1]. Immunotherapies have revolutionized the treatment of cancer due to their potential to elicit long-lasting cures, induce protection from recurrence, and impact distant metastatic disease [2–6]. The most successful immunotherapies for cancer include immune checkpoint blockade (ICB), engineered T cell therapies, and vaccines. However, ICB—the most commonly applied clinical immunotherapy—has only resulted in responses in only ∼12-33% of breast cancer patients [7]. Another commonly utilized strategy, engineered T cell therapies, has met with substantial struggles when tested in solid tumors, with only ∼33% of breast cancer patients in limited clinical trials demonstrating responses [8]. Further, cancer vaccines, which have been underutilized clinically, have only resulted in responses in ∼26-40% of melanoma patients [9–11] with little clinical application in breast cancer [12–14]. These therapies have struggled for many reasons, including tumor-mediated immune evasion, dysfunctional T cell activation, lack of appropriate cytokines, systemic pre-suppression due to prior treatment and continual tumor mutation [15–20]. Additionally, all of these therapies have focused on downstream effects of immunity, leaving the critical steps of antigen processing and presentation underexplored as therapeutic targets.

To improve immunotherapy responses, we first considered the various steps in the tumor immunity cycle [21]. In short, for an efficient immune response to develop against a cancer, cancer antigen must access an antigen-presenting cell and that antigen must then be presented to a cognate T cell alongside costimulatory molecules [21]. The cancer antigen-specific T cell then becomes activated, proliferates, and recirculates to reach the tumor, where it finds the correct antigen-bearing cancer cell and kills it [21].

The activation portions of this process typically occur in secondary lymphoid organs, such as lymph nodes (LNs), which are specialized organs that co-localize antigen, dendritic cells (DCs) and naïve T cells. The currently available therapeutic modalities (ICB, engineered T cells) focus on the T cell dynamics after (or without the need for) T cell activation, with few clinical approaches focused on targeted delivery or antigen presentation dynamics. As such, therapies targeting antigen processing and presentation are underexplored and have the potential to elicit substantial improvements in immunotherapy.

In this work, we demonstrate that breast cancer impairs the ability of tumor DCs or DCs in tumor-draining LNs (TdLNs) to efficiently present cancer neoantigen, which limits the potential to develop a productive anti-cancer immune response. We utilized LN-targeted adjuvant delivery strategies to induce antigen presentation and development of an antigen-specific T cell response, which resulted in long-term cures when combined with ICB. We also identified reduced IL1β levels within TdLNs as a potential cause for tumor-mediated impaired antigen presentation. We demonstrated that locoregional delivery of IL1β improves the capacity of DCs to present antigen and thus T cells to mount tumor antigen-specific responses. As a whole, this study demonstrates the potential of targeting antigen presentation in an antigen-agnostic manner to expand immunotherapy efficiency.

## RESULTS

### Tissue-resident DCs in tumor-draining LNs are impaired in their capacity to present neoantigens

The presentation of neoantigens is a critical step for developing and sustaining anti- cancer immune responses. Thus, we measured the ability of DCs from the spleen and LNs to present neoantigens in tumor-bearing and naïve animals. The LNs included in the analyses were the inguinal LN (Ing LN), which directly drains the fourth mammary fatpad (MFP) where the primary breast tumor was implanted, and the axillary LN (Ax LN), which is directly connected to the Ing LN via the inguinal-axillary lymphatic vessel. We evaluated LNs either ipsilateral to the tumor (Ipsi Ing LN and Ipsi Ax LN) or contralateral to the tumor (Contra Ing LN and Contra Ax LN) (Fig. 1A). To measure the ability of conventional DC-1s (cDC1, defined as CD45^+^CD3^-^CD11c^+^CD11b^-^CD8^+^Xcr1^+^), conventional DC-2s (cDC2, defined as CD45^+^CD3^-^CD11c^+^CD8^-^CD11b^+^major histocompatibility complex (MHC)II^+^), and plasmacytoid DCs (pDC, defined as CD45^+^CD3^-^CD11c^+^B220^+^) (Supplemental Figure 1) to process antigen, we implanted E0771 mammary carcinoma cells in the fourth MFP of immunocompetent C57/Bl6 mice. On day 7 of tumor growth, the relevant LNs (Fig. 1A), along with the spleen were excised and single cell suspensions were generated. Notably, this is a timeline in which metastatic burden is quite low in this model [22]. To evaluate antigen processing, we stimulated cells from animals bearing these tumors compared to tumor-naïve (naïve) animals ex vivo with DQ-OVA, a modified OVA protein in which the BODIPY fluorophore is self-quenched and emits no fluorescence in its native state but emits green fluorescence in its cleaved state, for 6 hours. Flow cytometry revealed that, as expected [23–25], cDC2s were more likely to process DQ-OVA when compared to cDC1s or pDCs (Fig. 1B). However, we saw no difference in the ability of each subset to process DQ-OVA based on tissue analyzed or whether it was from a tumor bearing or tumor naïve animal (Fig. 1B, Supplemental Figure 2A), indicating that the tumor had no effect on antigen processing.

**Fig. 1.**
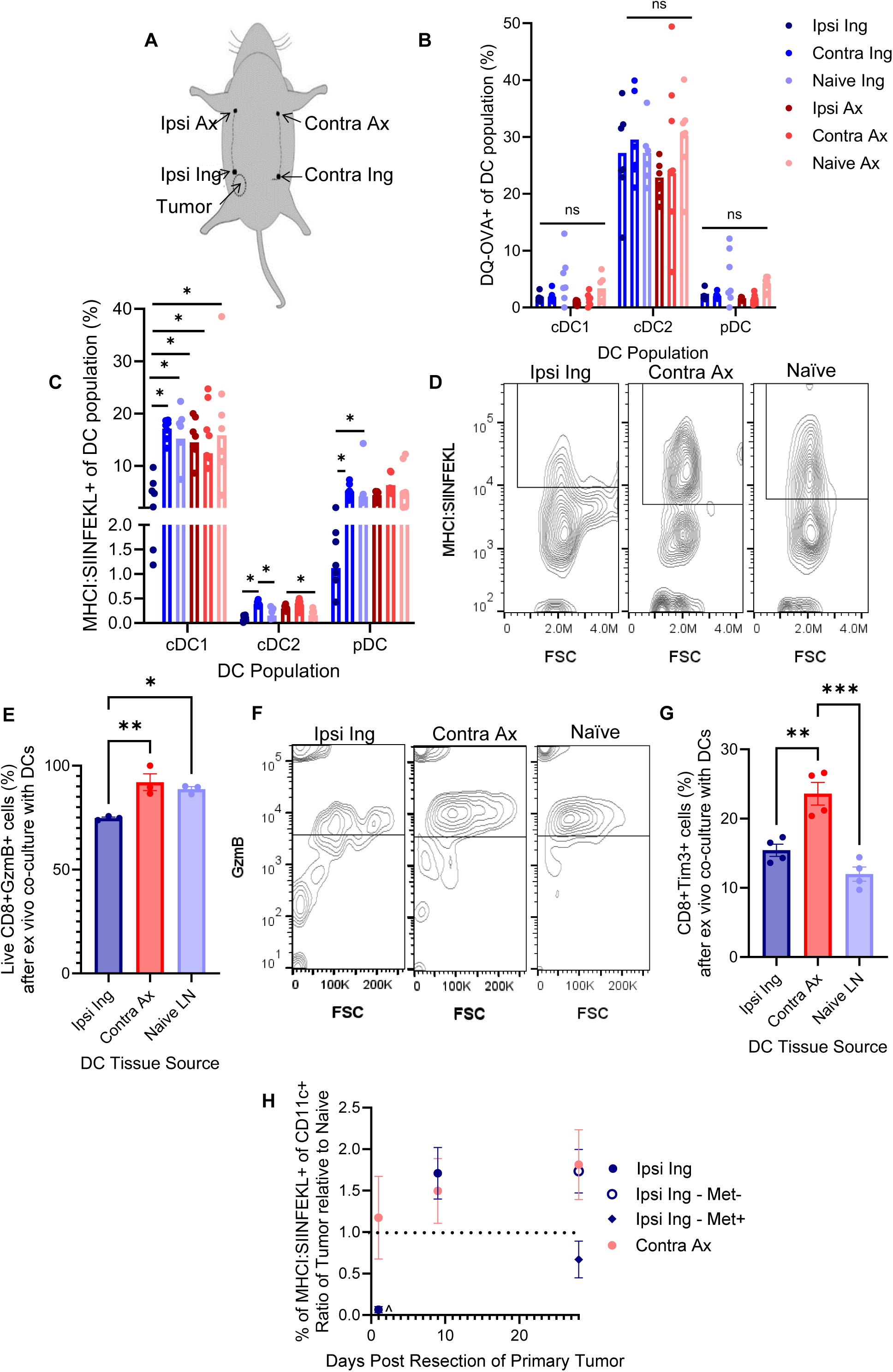
Locoregional LN tissue-resident cells are impaired in their capacity to present neoantigens. (**A**) Schematic of mouse indicating tissues of relevance. Frequency of DCs (cDC1 defined as CD11c^+^CD11b^-^CD8^+^Xcr1^+^; cDC2 defined as CD11c^+^CD8^-^CD11b^+^MHCII^+^; pDC defined as CD11c^+^B220^+^, Supplemental Figure 1) having processed DQ-OVA (**B**) or presenting SIINFEKL within the MHCI complex (as identified using the 25D1.16 mAb) (**C**) after cells from tissues of relevance from either naïve or day 7 E0771-bearing animals were co- incubated with DQ-OVA for 6 hours *ex vivo*. (**D**) Representative flow plots of MHCI:SIINFEKL staining (using 25D1.16 mAb) from cDC1s (defined above) derived from the Ipsi Ing, Contra Ax, or naïve LNs after co-incubation with DQ- OVA for 6 hours *ex vivo*. OT-I CD8^+^ T cells containing GzmB (Frequency, **E** representative flow plots, **F**) and Tim-3 (Frequency, **G**) after co-incubation with 100,000 DCs (by CD11c positive magnetic isolation) pre-incubated with OVA for 6 hours from different tissue sources, and then incubated with OT-I cells for 18 hours. (**H**) Ratio of frequency of DCs (total CD11c^+^) in various LNs presenting SIINFEKL within the MHCI complex after tumor resection in naïve relative to tumor-bearing animals (day 0 is the day of tumor resection which was preceded by 14 days of E0771 tumor growth). Met- indicates no macrometastatic disease; Met+ indicates macrometastatic disease per CD45^-^ staining. * indicates significance by two-way ANOVA with Tukey’s post-hoc test; ^ indicates difference against naïve animals by two-way ANOVA with Tukey’s post-hoc test; * or indicate p<0.05, ** indicates p<0.01; ns = not significant (p>0.05). n=3-8 animals.

We next assessed the ability of these DCs to present processed antigen using the 25D1.16 monoclonal antibody (mAb), which detects the OVA_257-264_ octapeptide (amino acid sequence: SIINFEKL) within the MHCI. This revealed that cDC1 were most likely to present SIINFEKL within the MHCI, followed by pDC, and cDC2 (Fig. 1C), consistent with the major functions of these distinct DC subsets [23–25]. When evaluating different DC subsets between these tissues, we found that cDC1s in the Ipsi Ing LN had markedly reduced presentation of SIINFEKL compared to cDC1s in any other LN analyzed (Fig. 1C-D), despite no difference in antigen processing (Fig. 1B). The lack of antigen presentation in the TdLN has potential ramifications on the development of an antigen-specific CD8^+^ T cell response. Among cDC1s within any LN not directly draining the tumors, we were not able to see any differences (Fig 1C-D), indicating that the impairment was locoregional. pDCs within the Ipsi Ing LN presented lower SIINFEKL within the MHCI in comparison to pDC within the Contra Ing and Naïve Ing LN, mimicking the trends seen among cDC1s in the Ipsi Ing LN most local to the tumor.

Finally, cDC2s had markedly lower presentation of SIINFEKL within the MHCI, as expected [23–25], but we did identify some differences in MHCI:SIINFEKL staining with cDC2s within the Ipsi Ing LN and Naïve Ing LN had lower SIINFEKL presentation compared to those in the Contra Ing LN. Further, cDC2s within the Ipsi Ax LN had lower SIINFEKL presentation compared to those in the Naïve Ax LN (Fig. 1C). Notably, the phenotype of DCs in these tissues was also altered by the presence of a tumor, with similar spatial dependence: DCs in the LN most proximal to the MFP tumor had the lowest levels of CD80, a marker of activation, and higher MHCI and MHCII levels (Supplemental Figure 3-4). Thus, distance relative to the primary tumor seemed to correlate with decreases in antigen presentation, with the Ipsi Ing LN (nearest the tumor) being most impaired. The Contra Ax LN farthest from the tumor was unaffected by the presence of a tumor (Supplemental Figure 2B). Overall, the impaired antigen presentation observed in the TdLNs (Fig. 1C-D) was not due to issues in antigen processing (Fig. 1B), but rather a deficiency in the mechanisms involved in antigen presentation. Examining the expression of genes involved in MHC recycling did not reveal any differences that could explain the differences in antigen presentation (Supplemental Fig. 5) [26–32], suggesting the impairment is not transcriptionally regulated.

In order to assess the functional ramifications of this impaired antigen presentation, we isolated CD11c^+^ cells from relevant tissues (the Ipsi Ing and Contra Ax LNs from E0771- bearing animals, compared to Ing and Ax LNs from naïve animals, Supplemental Figure 6). Due to concerns about the impacts of flow cytometry-associated sorting, we only evaluated CD11c^+^ cells isolated by magnetic sorting within 18 min, and not DC subsets. These cells were then incubated with OVA for 6 hours, washed, and then co-incubated with CD8^+^ T cells isolated from OT-I animals, which all have antigen-specificity towards the SIINFEKL antigen. OT-I cells were then stained for phenotypic changes. This revealed that OT-I CD8^+^ T cells incubated with DCs isolated from the Ipsi Ing LN, which directly drains the tumor, had lower granzyme B (GzmB) and TIM-3 expression (Fig. 1E- F) when compared to OT-I CD8^+^ T cells incubated with DCs from the Contra Ax or naïve LNs, indicating lower functional CD8^+^ T cell cytotoxicity. These data demonstrate that the deficits we measured in the ability of DCs to present antigen in TdLNs have functional ramifications for the ability of the animal to activate antigen-specific CD8^+^ T cells.

Finally, we assessed the persistence of the locoregional LN antigen presentation impairment after tumor removal. To this end, we orthotopically implanted immunocompetent C57/Bl6 mice with E0771 tumors. On day 14 of tumor growth, tumors were resected. Animals were sacrificed 1 day (to assess immediate impacts of tumor removal), 9 days (to assess a timepoint in which acute tumor-mediated effects could be ameliorated) or 28 days (to assess potentially metastatic lesions) later and antigen presentation was assessed by flow cytometry. On day 1 after tumor resection, antigen presentation impairment was similar to that seen in animals with intact tumors (Fig. 1G, Supplemental Fig. 7). On day 9 post tumor resection, antigen presentation was improved, and was similar in the Ipsi Ing LN compared to the Contra Ax LN and LNs from naïve animals (Fig. 1G, Supplemental Fig. 7-8). On day 28 after tumor resection, antigen presentation was impaired in LNs with metastatic lesions, but in TdLNs without metastasis, antigen presentation was similar to distant LNs in (Fig. 1G, Supplemental Fig. 7-8). Thus, the presence of cancer or cancer-disseminated factors interferes with antigen presentation.

### Phenotype of TdLN-resident DCs is altered

To test the hypothesis that tumor-disseminated factors were responsible for the alterations in tumor antigen presentation, we performed multiplexed cytokine assays on tissues (tumors or naïve MFP, Ipsi Ing LNs, and Contra Ax LNs) derived from tumor-bearing or naïve animals (Fig. 2A, Supplemental Fig. 9). This revealed differences in IL1β (total, includes both pro and active forms) in the Ipsi Ing LN in tumor-bearing versus naïve animals (Fig. 2A, Supplemental Fig. 9), but not the Contra Ax LN. To test the effects of these cytokines, we measured differences in antigen presentation after ex vivo culture of cells from tumor-naïve animals in the presence of IL1β, IL33, MCP or IP10 with OVA (Fig. 2B, Supplemental Fig. 9). The only clear difference in antigen presentation was with the addition of IL1β, in which antigen presentation was enhanced (Fig. 2B, Supplemental Fig. 9). As IL1β concentration was decreased in the tumor and Ipsi Ing LN of tumor-bearing animals compared to naïve animals, but unchanged in the Contra Ax LN (Fig. 2A), we hypothesized that changes in IL1β levels could explain the impaired DC antigen presentation (Fig. 1). As such, we isolated CD11c^+^ cells from tissues of relevance from tumor-bearing versus naïve animals. These cells were then incubated with OVA in the presence of IL-1β or vehicle (saline) for 6 hours, washed, co-cultured with CD8^+^ T cells isolated from OT-I animals for 18 hours, and then phenotype of T cells was measured using flow cytometry. This revealed that the addition of IL1β enhanced GzmB production in both the Ipsi Ing and Contra Ax LNs CD11c+ cells from tumor bearing animals (Fig. 2C-D, Supplemental Figure 10), without differences in Tcf1 or programmed death-1 (PD1) expression (Supplemental Figure 9), indicating that the addition of IL1β in tumor-bearing animals has the potential to induce a functional anti-tumor T cell response.

**Fig. 2.**
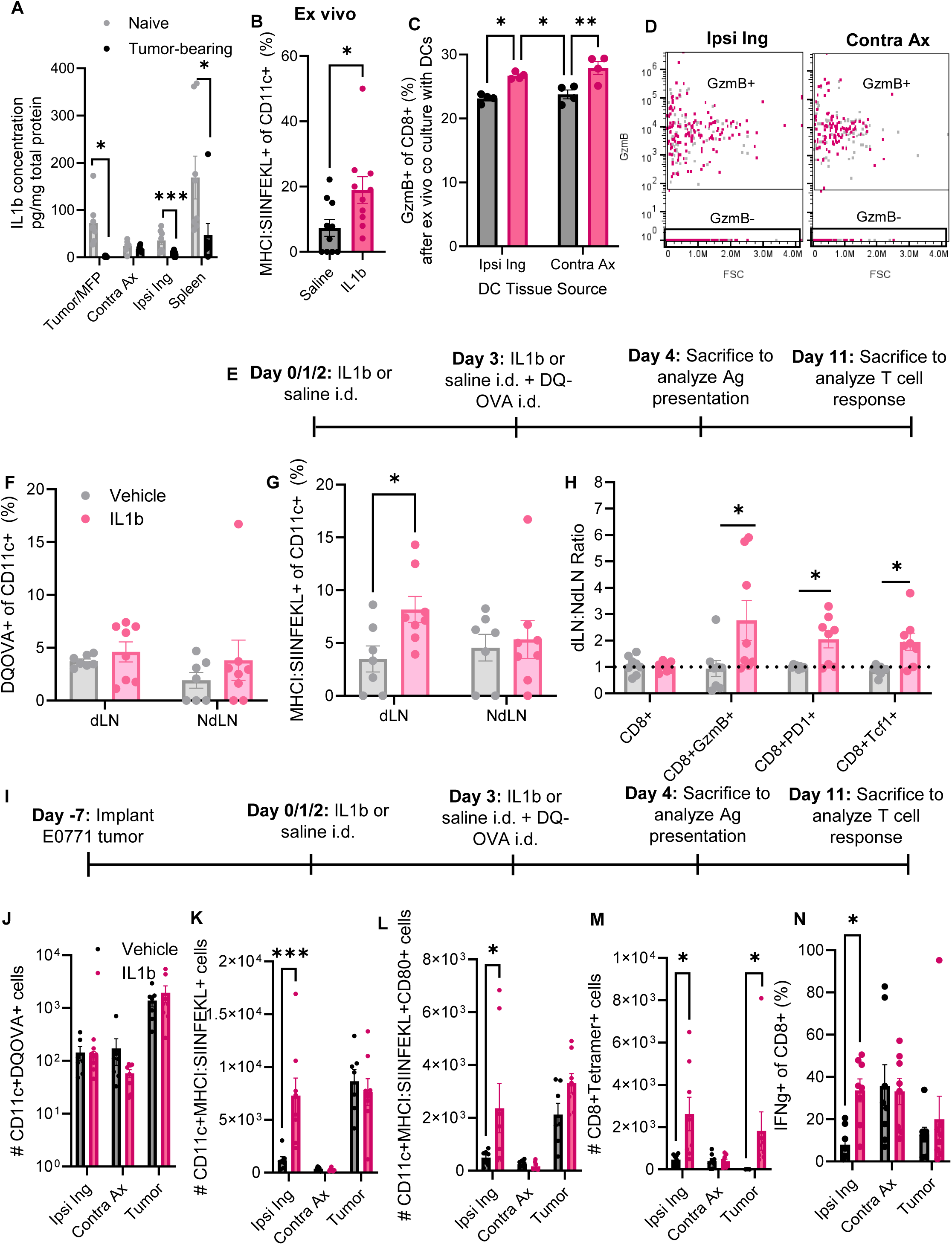
Targeted delivery of IL1β enhances antigen presentation and alters T cell response in both naïve and tumor-bearing animals. (**A**) IL1β levels in relevant tissues of day 14 E0771-bearing animals as measured by ELISA. (**B**) Presentation of SIINFEKL antigen within the MHCI complex after co-incubation with DQ-OVA for 6 hours ex vivo with or without IL1β. Frequency (**C**) and representative flow plots (**D**) of GzmB within OT-I CD8^+^ (by CD8 negative magnetic isolation) T cells after 6 hours of co-incubation (Brefeldin-A added at 3 hours) with 100,000 DCs (by CD11c positive magnetic isolation) with or without IL1β (1ng). (**E**) Timeline for experiment evaluating effects of IL1β (1ng at each dose) on antigen presentation and development of an antigen-specific T cell response in naïve animals. Frequency of CD11c^+^ cells processing DQ-OVA (**F**) or presenting SIINFEKL within the MHCI complex (**G**) after 6 hour incubation with DQ-OVA, from tissues from animals treated based on configuration in 2E. (**H**) Endogenous T cell phenotype in dLNs IL1β treatment relative to NdLNs IL1β treatment in tumor-naïve animals. (**I**) Timeline for experiment evaluating role of IL1β (1ng at each dose) on antigen presentation and development of an antigen- specific T cell response in tumor-bearing animals. Number of CD11c^+^ cells processing antigen (as measured by cleaved DQ-OVA), **J**), presenting SIINFEKL within the MHCI complex (**K**), and both presenting SIINFEKL within the MHCI and stained with CD80 (**L**) after 6 hour ex vivo incubation with DQ-OVA, from tissues from animals treated based on the configuration in 2I. (**M**) Number of SIINFEKL tetramer-specific CD8^+^ T cells after experiment defined in 2I. (**N**) Frequency of CD8^+^ T cells stained for IFN (interferon)-γ after experiment defined in 2I. * indicates significance by one-way ANOVA with Tukey’s post-hoc test (A, C, G-H, J-N), t-test (B, D). * indicates p<0.05, ** indicates p<0.01, *** indicates p<0.005, n=6-8 animals.

As in vitro cultures do not completely recapitulate the complexity of the immune system and immune reactions, we next tested the effects of IL1β on in vivo antigen presentation in tumor-naïve animals. Our data revealed that IL1β had no impact on antigen processing in DCs as assessed by DQ-OVA positivity (Fig. 2E-F, Supplemental Fig. 11). However, the addition of IL1β enhanced antigen presentation measured by the frequency of CD11c^+^MHCI:SIINFEKL^+^ DCs in the LN draining the IL1β injection (dLN, Fig. 2G, Supplemental Fig. 11). We next assessed the impacts of these changes in DC antigen presentation on T cell responses. The dLN the IL1β injection showed no change in the number of total endogenous CD8^+^ T cells compared to vehicle administration or to distant LNs in the same animal (non-draining LN, NdLN) (Fig. 2H). However, GzmB+, PD1+ or Tcf1+ CD8^+^ T cells were enhanced in LNs that received IL1β compared to animals receiving vehicle control (Fig. 2H). Thus, in cancer naïve animals, IL1β was able to enhance antigen presentation and T cell activation compared to vehicle controls.

We next assessed the impact of IL1β injection targeting the TdLN in a cancer context in vivo (Fig. 2I). IL1β injection into the locoregional flank skin did not change the number of DCs taking up and processing the antigen as assessed by DQ-OVA (Fig. 2J, Supplemental Fig. 11) but did increase the number of DCs presenting antigen (MHCI:SIINFEKL^+^) within the Ipsi Ing LN compared to mice that received vehicle control (Fig. 2K, Supplemental Fig. 11). There were also more CD80^+^ cells presenting SIINFEKL after IL1β delivery in the Ipsi Ing LN, indicating more reactivity by these DCs (Fig. 2L). Finally, we measured the development of antigen-specific T cells using SIINFEKL-specific tetramers. The data show that there were more tumor antigen- specific T cells in both the Ipsi Ing LN and tumor of animals that received IL1β compared to those that received vehicle control (Fig. 2M, Supplemental Fig. 11).

Production of interferon (IFN)-γ by CD8^+^ T cells in LNs treated with IL1β was also enhanced in the Ipsi Ing LN with delivery of IL1β (Fig. 2N), indicating T cell functionality. Thus, IL1β was able to improve the antigen presentation by DCs and generate a T cell response, potentially overcoming the antigen presentation impairment identified in Fig. 1.

### Delivering immunological adjuvant to distant LNs enhances antigen presentation and antigen-specific T cells

Given that antigen presentation was impaired in locoregional LNs, we next tried a strategy aimed at initiating immunity in distant LNs [33], thus bypassing the immunosuppression that exists near the tumor. We therefore hypothesized that introduction of antigen and adjuvant to these distant, non-suppressed nodes would enhance antigen presentation and T cell activation. To determine the appropriate location for injection, we first mapped the distribution of a surrogate antigen (FITC- conjugated 2 MDa dextran) after injection in either the ipsilateral flank skin or contralateral forelimb (Fig. 3A). After injection into the ipsilateral flank skin, surrogate antigen accumulated in the Ipsi Ing LN and Ipsi Ax LN (which drains the Ipsi Ing LN) at approximately 10^2^-10^3^ fold higher rates compared to the Contra LNs (Fig. 3B-C). When surrogate antigen was injected in the contralateral forelimb, it accumulated in the Contra Ax LN (draining the injection site) at approximately 10-fold higher rates compared to any other LN (Fig. 3C). Thus, these injection sites could be used to target specific LNs.

**Fig. 3.**
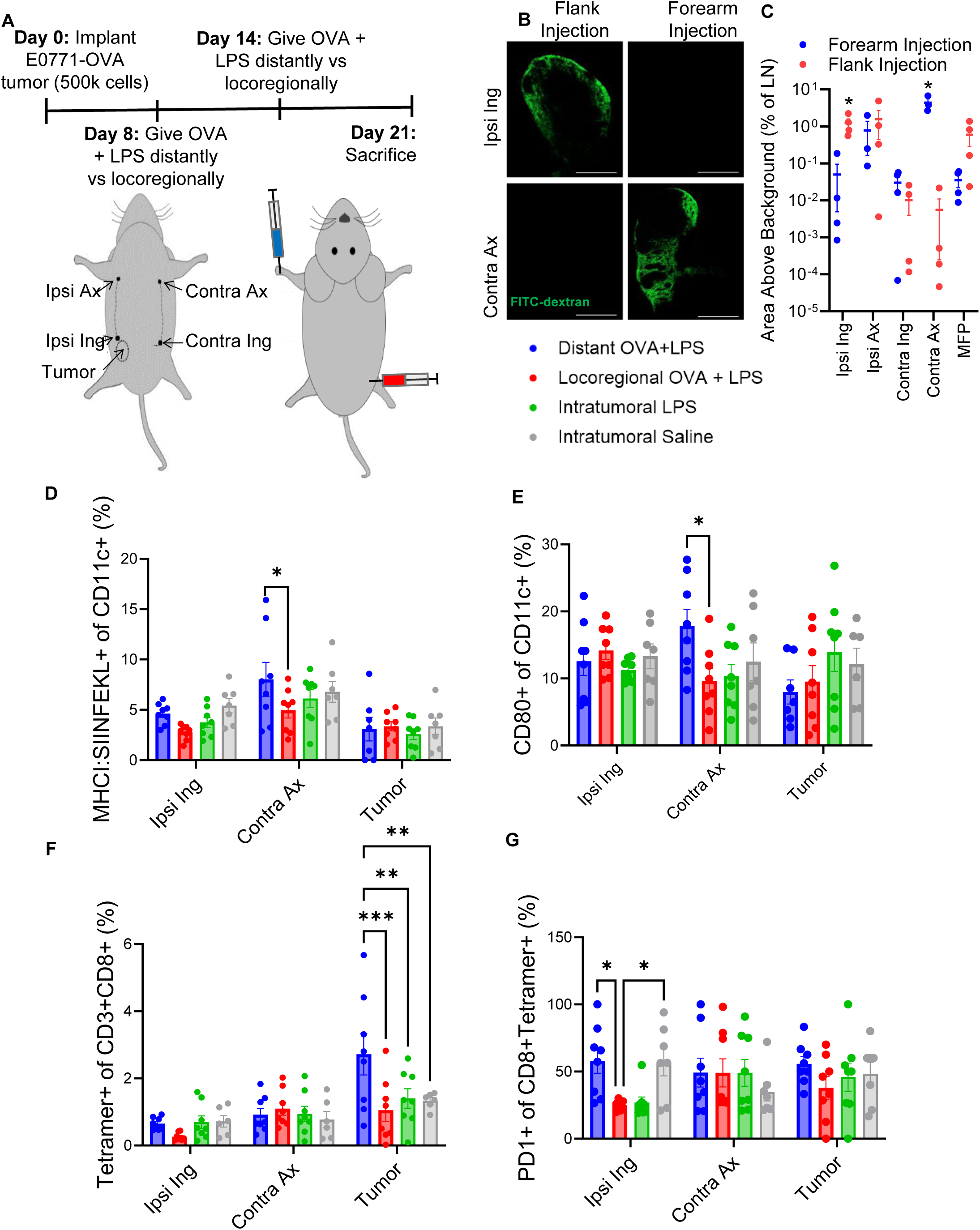
Delivering immunological adjuvant to distant LNs enhances antigen presentation and antigen-specific T cells. (**A**) Experimental design and schema showing relevant tissues, and injection schemes in mice for experiments evaluating effects of location of therapeutic immunological adjuvant on antigen presentation and development of antigen-specific T cell responses. (**B**) Images demonstrating accumulation of 2MDa, fluorescently tagged dextran in LNs after injection into the flank or forelimb. (**C**) Area above background (based on animal injected with saline) within LNs after injection of fluorescent dextran into flank or forelimb. (**D**) Frequency of CD11c^+^ cells with SIINFEKL loaded in the MHCI complex, within relevant tissues after delivery of immunological adjuvant. (**E**) Frequency of CD11c^+^ cells expressing CD80 after delivery of immunological adjuvant. (**F**) Frequency of CD8^+^ T cells tagged with SIINFEKL-specific tetramer in relevant tissues, (**G**) PD1^+^ T cells of tetramer specific T cells in relevant tissues. * indicates significance by two-way ANOVA with Tukey’s post-hoc test. * indicates p<0.05; ** indicates p<0.01; *** indicates p<0.005. n=7 (saline group) – 8 (treated groups) animals. Scale bar = 500µm

To test the impact of an altered delivery strategy on response to immunization, E0771- OVA tumors were implanted into C57/Bl6 mice, and OVA antigen in combination with lipopolysaccharide (LPS), an immunological adjuvant commonly used preclinically, (OVA+LPS) were injected into either the ipsilateral flank (Locoregional OVA+LPS) or contralateral forelimb (Distant OVA+LPS) on day 8 of tumor growth followed by a boost on day 14 (Fig. 3A). When the LPS+OVA combination was injected in the ipsilateral flank, there were few differences in antigen presentation by CD11c^+^ cells in the targeted LN (the Ipsi Ing LN), the non-targeted LN (Contra Ax), or the tumor (Fig. 3D, Supplemental Fig. 12). When the LPS+OVA were injected in the contralateral forelimb, antigen presentation by CD11c^+^ cells was enhanced in the targeted LN (the Contra Ax LN), but not the non-targeted LN (Ipsi Ing LN) or the tumor (Fig. 3D, Supplemental Fig. 12). Neither intratumoral (i.t.) delivery of OVA+LPS or vehicle altered antigen presentation by CD11c^+^ cells (Fig. 3D, Supplemental Fig. 12). When comparing across groups, only the distant OVA+LPS induced an increase in antigen presentation in any tissue when compared to the saline group (Fig. 3D, Supplemental Fig. 12). We also observed a very similar trend when examining the DC activation marker CD80 in these tissues, with only DCs in the Contra Ax LN demonstrating an enhancement in CD80 after forelimb delivery of OVA+LPS (Fig. 3E). As a whole, these data indicate that targeting LNs that do not directly drain the tumor has the potential to enhance both quantity and quality of antigen presentation.

We next assessed the impacts of different delivery strategies on the development of an antigen-specific response using SIINFEKL tetramers to identify antigen-specific T cell activation. When OVA+LPS was delivered targeting a distant LN, there was no change in the frequency of antigen-specific T cells in the Ipsi Ing or Contra Ax LNs compared to the saline group, but distant LN-targeted OVA+LPS (Distant OVA+LPS) did increase infiltration of antigen-specific T cells into the tumor when compared to the saline group (Fig. 3F, Supplemental Fig. 12). Likewise, the frequency of SIINFEKL-specific T cells expressing PD1 increased in the Ipsi Ing LN when treated with Distant OVA+LPS compared to locoregional or i.t. OVA+LPS, indicating enhanced antigen experience of those cells (Fig. 3G). When OVA+LPS was delivered locoregionally or i.t., no changes in antigen-specific T cells were observed in the Ipsi Ing or Contra Ax LNs or in the tumor when compared to the saline group (Fig. 3F, Supplemental Fig. 12). Thus, only distant OVA+LPS was able to enhance infiltration of antigen-specific T cells into the tumor, a marker that has been shown to positively correlate with survival [34–38]. Interestingly, the Ipsi Ax LN, which directly drains the Ipsi Ing LN via the inguinal-axillary lymphatic vessel, showed very similar trends in the presence of antigen-specific T cells, indicating that improvements after locoregional injections could persist beyond the very first TdLN into a secondary LN (Supplemental Figure 12). Because immunological adjuvants are typically utilized in vaccines to increase the development of antibody responses, we measured anti-OVA IgG in serum of these animals, which did not reveal differences in the development of anti-OVA antibodies based on the location of injection (though a clear difference was noted based on the formulation of injection) (Supplemental Fig. 13).

### Delivering immunological adjuvant to distant LNs delays tumor growth

Strategies for combining antigen with immunological adjuvant require the availability and knowledge of a tumor-associated antigen. In the clinic, this is a challenge because of the wide variety of mutations leading to the development of a tumor and continued tumor mutation throughout disease progression [39–42]. Because of this limitation, we next tested the ability of two clinically relevant immunological adjuvants, aluminum hydroxide salt (alum) used in TDaP (tetanus, diphtheria, and pertussis), hepatitis, and human papillomavirus vaccines; and Titermax Gold (Titermax), an analog to MF59 used in influenza vaccines [43–46], to slow tumor growth in two breast tumor models (4T1 and E0771) without addition of exogenous antigen, i.e. in an antigen-agnostic manner (Fig. 4A). Delivery of either adjuvant slowed tumor growth and enhanced survival when delivered distantly from the tumor, but not locoregionally or i.t. (Fig. 4B-C, Supplemental Fig. 14-15). These cohorts were sacrificed on day 28 of tumor growth to analyze the infiltration of cells into the tumor. This revealed that distant delivery of immunological adjuvant increased the infiltration of CD8^+^ T cells into the tumor (Fig. 4D, Supplemental Fig. 14) and with a higher proportion of GzmB^+^ cytotoxic CD8^+^ cells into the tumor (Fig. 4E, Supplemental Fig. 14). There were no differences in PD1^+^ or Tcf1^+^ infiltrates (Supplemental Fig. 14). Thus, distant delivery of immunological adjuvant alone (i.e. antigen-agnostically) was able to slow tumor growth in two different tumor models and enhance animal survival, independent of the type of immunological adjuvant itself, and in an antigen-agnostic manner.

**Fig. 4.**
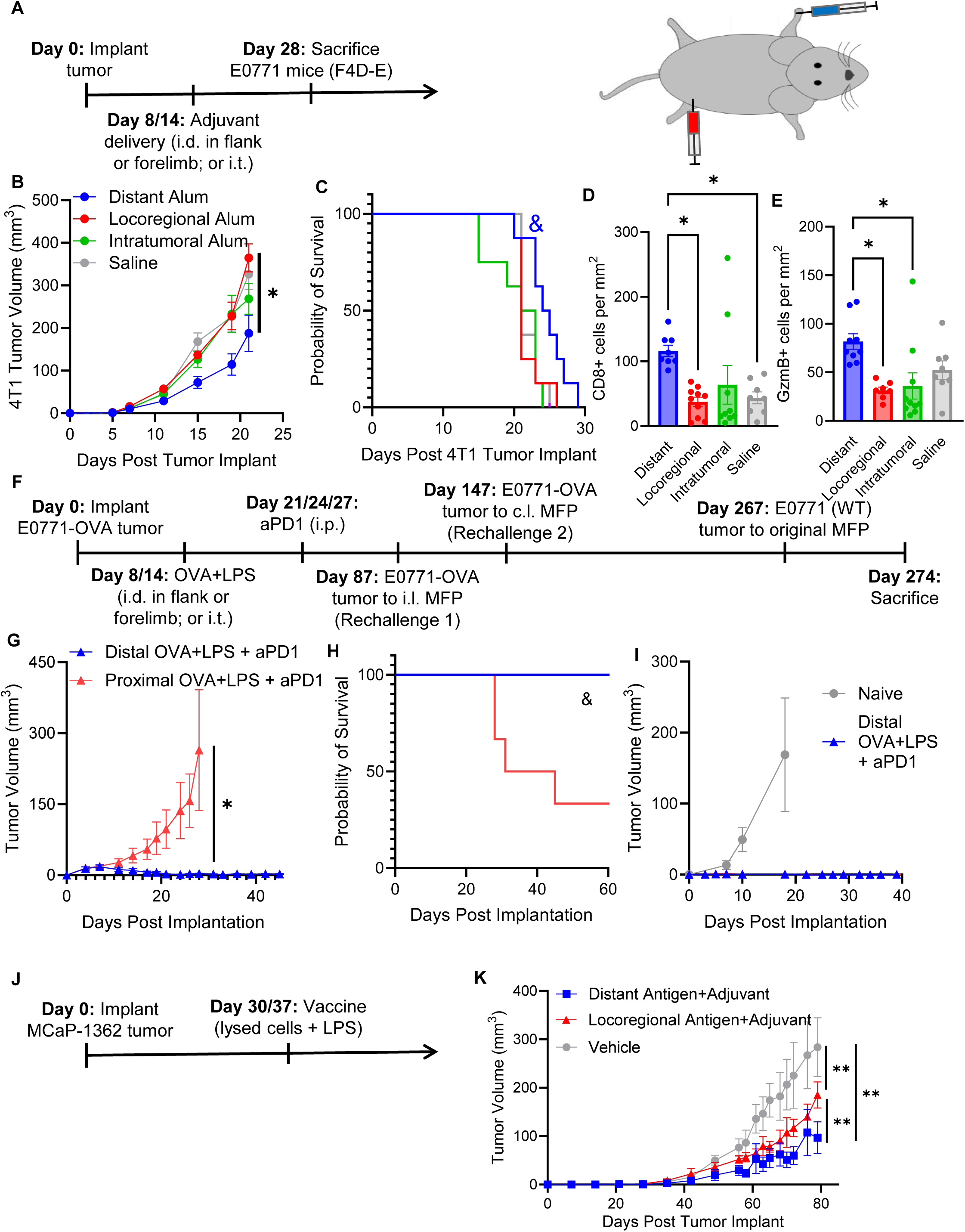
Delivering adjuvant to distant LNs slows tumor growth. (**A**) General timeline for experiments investigating impacts of adjuvant delivery on tumor growth dynamics, and schematic of injections into mouse. Tumor growth (**B**) and survival (**C**) after alum was delivered i.d. distant, locoregionally, or i.t. to 4T1-bearing animals compared to growth after i.t. saline delivery. Number of CD8^+^ (**D**) and GzmB^+^ (**E**) cells per mm^2^ in sections of 4T1 tumors after i.d. distant or locoregional, or i.t. Titermax, or vehicle delivery. **(F**) Experimental timeline for combination of delivery of distant versus local adjuvant with ICB (αPD1) and rechallenge for surviving animals. Tumor growth (**G**) and survival (**H**) of E0771- bearing animals treated with distant or locoregional adjuvant (LPS+OVA) and ICB (αPD1) or isotype IgG. All surviving animals were maintained at least 100 days after no evidence of disease was established. (**I**) Tumor growth after E0771-OVA cells were implanted in the original tumor location of surviving animals compared to tumor-naïve animals implanted on the same day. Experimental design (**J**) and tumor growth (**K**) after MCaP-bearing animals were treated locoregionally or distantly with lysed tumor cells combined with LPS, compared to i.t. saline. * indicates significance by repeat measures (RM) one-way ANOVA (B, G, K) or one-way ANOVA (D-E); & indicates significance relative to all other groups by log-rank test. * and & indicate p<0.05, ** indicates p<0.01.

As these antigen-agnostic strategies did not result in curative responses, we sought to combine immunological adjuvant delivery with ICB in the form of systemic αPD1 mAb delivery. Mice bearing E0771-OVA tumors were vaccinated with LPS+OVA on day 8, followed by a boost on day 14 (Fig. 4F, Supplemental Figure 15). On day 21, a timepoint in which antigen-specific T cells infiltrated the tumor to a higher degree after distant OVA+LPS treatment (Fig. 3), the first of 3 doses of αPD1 mAb or isotype IgG was given intraperitoneally. Slower tumor growth in animals receiving the combined distant adjuvant and αPD1 was measured when compared to those receiving locoregional adjuvant and αPD1 (Fig. 4G, Supplemental Fig. 16-17). Distant OVA+LPS with αPD1 induced long-term survival in 100% of these animals (Fig. 4H, Supplemental Fig. 16-17), with protection from three different rechallenges (Fig. 4I, Supplemental Fig. 16-17), demonstrating an effective and long-lasting anti-tumor immune response.

Finally, we investigated the capacity of animals implanted with the MCaP-1362 breast carcinoma line to respond to a “vaccine” comprised of lysed MCaP-1362 cells combined with LPS when delivered locoregionally versus distantly (Fig. 4J), in order to understand the effects of a less immunogenic and more physiological antigen source. This revealed that delivery of locoregional antigen+adjuvant slowed tumor growth, but distant delivery of tumor antigen+adjuvant further slowed tumor growth (Fig. 4K). These data indicate that even in a clinically-relevant situation in which a lysed tumor sample is used, distant delivery could be beneficial. Thus, overall, combining distant delivery of immunological adjuvant with ICB reduces tumor burden, enhances animal survival, and induces the development of a robust and long-term, curative immunological memory response.

### IL1**β** in combination with other immunotherapies, but not as a monotherapy, slows tumor growth

We noted that distant adjuvant delivery induced higher IL1β within the tumor, which may be responsible for the effects we saw in antigen presentation, antigen-specific T cell development, and tumor growth dynamics with distant adjuvant delivery (Supplemental Fig. 18). As such, we sought to explore locoregional IL1β delivery as an alternative therapeutic option. As our data showed that IL1β delivery to LNs could improve antigen presentation by DCs, we tested the hypothesis that IL1β supplementation could improve tumor responses. In order to measure the capacity of IL1β to impact tumor growth, we orthotopically implanted animals with E0771 tumors, and delivered IL1β as a monotherapy i.t. (Fig. 5A). IL1β monotherapy showed only a very slight slowing of tumor growth but did reach significance in overall survival (Fig. 5B, Supplemental Fig. 19). However, only 1 of 7 animals survived long-term (Fig. 5B), which is not sufficient.

**Fig. 5.**
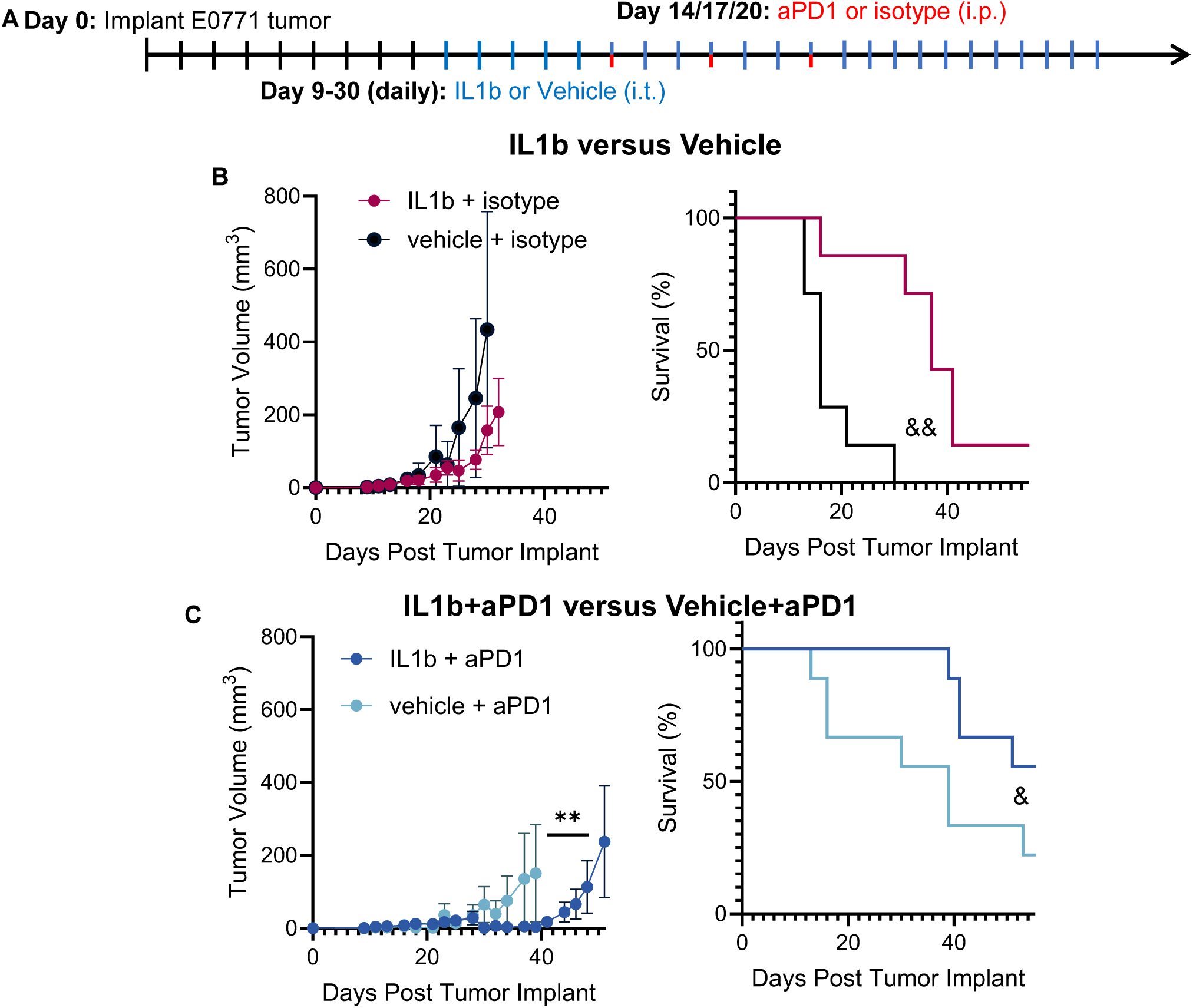
Local IL1β when combined with systemic aPD1 induces long-term responses against breast cancer. (**A**) Timeline for survival experiment study when IL1β (1ng at each dosing) or vehicle (saline) is combined with αPD1 or isotype mAb in E0771 tumor-bearing animals. Tumor growth until humane endpoint reached in first animal in cohort (left), and survival (right) after delivery of IL1β (1ng at each dosing) or vehicle alone in combination with isotype mAb (**B**), or αPD1 (**C**). All surviving animals were maintained at least 100 days after no evidence of disease was established. n=7-9 animals. * indicates significance by RM-ANOVA; & indicates significance by log-rank test against all other groups. & indicates p<0.05, ** or && indicate p<0.01.

As our data showed that IL1β enhanced DC antigen presentation and subsequent T cell activation, we hypothesized that tumor-mediated impairment of T cell effector function could be preventing ultimate anti-cancer effector immune responses that would improve survival. Thus, we combined IL1β delivery with ICB (αPD-1 mAb) to maintain functionality of effector T cells after IL1β assisted activation (Fig. 5A). The data show a slowing of tumor growth in animals treated with this combination (Fig. 5C, Supplemental Fig. 19). Survival curves indicated some variability in long term survival in the IL1β+αPD1 group but indicate that this combination improved survival compared to both monotherapies (Fig. 5C, Supplemental Fig. 19). Notably, the αPD1 alone group performed worse than expected in this experiment [33,47], so this is a substantial improvement compared to expectations. All animals surviving the original tumor implantation (treated in both the IL1β monotherapy and IL1β+αPD1 therapy groups) survived tumor rechallenge with 2x10^6^ E0771 cells with no evidence of disease at any point, indicating development of a durable immune response. Thus, combining IL1β with αPD1 induced an improved response in a subset of animals, and has the potential to improve clinical outcomes.

## DISCUSSION

Our data show that breast tumors impair antigen presentation in locoregional LNs, the effects of which can be compensated via either IL1β delivery to the TdLN or via targeted adjuvant delivery strategies to distant LNs not suppressed by the tumor. This simple, targeted injection strategy could be easily applied in the clinic to improve upon current rates of durable responses to ICB therapy in a variety of patient populations. Here we showed how tumors can affect this process, specifically by affecting MHCI antigen presentation in TdLNs and on CD8^+^ T cells. The impacts of tumor-mediated factors on MHCII antigen presentation and CD4^+^ T cell responses are subjects for future research efforts.

We identified reduced IL1β as a key barrier to the process of antigen presentation by DCs in TdLNs and showed that i.t. delivery of IL1β in combination with αPD1 enhanced survival and slowed tumor growth. As cytokine therapies have traditionally had limited translatability, to overcome potential translational issues with an IL1β centered strategy, we utilized specific antigen-agnostic adjuvant delivery strategies to specific LNs that access tumor-disseminated antigen, but are not as immunosuppressed in the way direct TdLNs are (Jeanbart et al., 2014). The potential for translation of this technique has great promise due to its simplicity and the use of immunological adjuvant without the need for understanding the changing neoantigens within the cancer itself. However, the differences between preclinical and clinical situations may result in problems in translation. The mouse lymphatic network is much simpler when compared to the human lymphatic network. For example, in a mouse, there is typically one Ax LN on either side of the mouse, while in the human, there can be up to dozens of interconnected LNs [49–52]. Likewise, the length-scales of these interactions differ substantially. We utilized the most distant LN that was in the same general lymphatic network in these studies, but acknowledge that the length-scales in mice is still not directly comparable to humans. These differences make the targeting of the specific LN of interest more challenging in the clinic. Computational approaches can be utilized to understand ideal LNs to target for an individual patient [53] and improve chances of durable responses even in late-stage breast cancer patients. Notably, we only utilize preclinical models of breast cancer in this study, most experiments of which are done in the E0771 model of luminal breast cancer (with some experiments occurring in the 4T1 triple-negative breast cancer and MCaP-1362 breast cancer line). However, we nonetheless hypothesize that these results could impact patients with other types of cancer, as the majority of our findings are immune cell-intrinsic, not cancer cell-intrinsic, implying potential commonalities and implications for additional patient populations.

Overall, this work demonstrates that locoregional antigen presentation is impaired in a tumor context, a process mediated by IL1β. This in turn impairs the development of an anti-cancer T cell response, which allows the tumor to progress. However, by targeting immunological adjuvant to a distant LN or locoregionally delivering IL1β, we demonstrate prolonged survival. This work has potential implications for therapeutic interventions and shines light on an underexplored and exciting mechanism by which to target the tumor immunity cycle.

## MATERIALS AND METHODS

### Sex as a Biological Variable

Only female animals were included in this study, as the study focuses on breast cancer, which presents quite differently in males compared to females.

### Animals and Tumor Models

We used three different syngeneic breast tumor lines in this work. E0771 mammary carcinoma cells (ATCC), MCaP-1362 mammary carcinoma cells (developed within Edwin L. Steele Laboratories), and 4T1 mammary carcinoma (ATCC), were grown in vitro in Dulbecco’s Modified Eagle’s Medium (Gibco, Invitrogen Life Technologies) containing 10% fetal bovine serum (FBS, Atlanta Biologicals) and 1% penicillin-streptomycin (Gibco, Invitrogen Life Technologies). Cells were maintained in a 5% CO_2_, humidified incubator at 37°C. All cells were recently authenticated and mycoplasma free. E0771 cells (0.5 x 10^6^), MCaP cells (10^6^), or 4T1 cells (0.5 x 10^6^) in saline (total volume 30µL) were implanted in the fourth MFP of female immunocompetent C57/Bl6 (E0771) or BALB/c (4T1 and MCaP) mice (aged 6-10 weeks). Animals were monitored every 1-7 day throughout tumor development and progression. For any survival experiments, surviving animals were maintained at least 100 days without evidence of disease. All procedures were performed according to the guidelines of the Institutional Animal Care and Use Committee of the Massachusetts General Hospital (Protocol #2011N000085, 2004N000123).

### Flow cytometry

Animals were sacrificed in accordance with protocol guidelines. Tissues were resected and maintained on ice in Hank’s Buffered Salt Solution (HBSS). Tissues were processed by mechanical dissociation through a 70 μm cell strainer and washed with HBSS. For spleen samples, cells were resuspended in ACK lysis buffer (Thermo Fisher) for 7 min at room temperature and then quenched with approximately 30mL phosphate-buffered saline (PBS). Samples were spun down for 5 min at 350g and plated in a 96-well U-bottom plate. The plate(s) were spun down for 5 min at 350g and Zombie UV/NIR/Red (based on experiment) Live/Dead (1:100, Biolegend) was added to samples, and allowed to incubate for 30 min at room temperature in the dark. Samples were spun down for 5 min at 350g and washed with PBS. Fc blocking antibody (anti-CD16/32 1:200, Tonbo Bioscience) was added to samples and allowed to incubate for 30 min at room temperature in the dark. Samples were spun down for 5 min at 350g and surface stain antibodies (Supplemental Table 1) were added. Samples were allowed to incubate for 30 min on ice in the dark. Samples were spun down for 5 min at 350g and washed with PBS. For experiments without intracellular stains, fixative/permeabilization buffer (eBioscience) was added to each sample and allowed to incubate for 30 minutes on ice in the dark. Samples were spun down for 5 min at 350g, washed with eBioscience buffer, and spun down again. Intracellular stains (Supplemental Table 1) were then added and samples allowed to incubate for 30 min on ice in the dark. Samples were spun down for 5 min at 350g, washed with eBioscience buffer, and spun down again. FACS buffer (1% FBS in PBS) was added to each sample, and then samples were spun down for 5 min at 350g, washed, and maintained at 4°C until analyzed via a customized BD LSRII or Cytek Aurora flow cytometer. Data were then analyzed using FlowJo version 10.

### Ex vivo stimulation

Animals were sacrificed in accordance with MGH guidelines and tissues excised. Tissues were mechanically dissociated through 70 μm filters to generate single cell suspensions. For spleen and blood samples, cells were resuspended in ACK Lysis buffer (Thermo Fisher) and allowed to incubate for 7 min at room temperature. Samples were then quenched with ∼30mL PBS and spun down for 5 min at 350g. Cells were plated in 96-well U-bottom plates, spun down for 5 min at 350g, and resuspended in 100 µL of appropriate media (Roswell Park Memorial Institute media, RPMI (Gibco, Invitrogen Life Sciences) + 10% FBS for unstimulated samples;

RPMI (Gibco, Invitrogen Life Sciences) + 10% FBS + 10 µg OVA (Sigma Aldrich) for OVA-stimulated samples; RPMI (Gibco, Invitrogen Life Sciences + 10% FBS + 10 µg DQ-OVA (Thermo Fisher) for DQ-OVA-stimulated samples; RPMI (Gibco, Invitrogen Life Sciences) + 10% FBS + 1 ng cytokine (Recombinant IL1β, IL44, MCP, or IP10; Biolegend) + 10 µg OVA (Sigma Aldrich) for cytokine experiments) and allowed to incubate for 6 hours in a 5% CO_2_, humidified incubator at 37°C. Cells were then stained for flow cytometry as above.

### Tumor resection

Animals were given pre-procedure analgesic (0.1 mg/kg buprenorphine) and anesthetized using ketamine-xylazine (80-100 mg/kg ketamine + 5- 10 mg/kg xylazine). Eyes were lubricated with Lubricant PM ointment (AACE Pharmaceuticals). Animals were then placed on a warming bench, and the surgical site prepared. Any remaining hair around the surgical site was shaved, and then the site was cleaned alternatively between betadine scrub and alcohol a total of three times.

Once animals were prepared, an incision was made around the tumor site, and blunt dissection was used to separate the tumor from underlying fascia. The tumor was then resected and any bleeding controlled via pressure. The skin was then closed with wound clips. Post-procedure analgesic (0.1 mg/kg buprenorphine) was maintained for 72 hours after termination of the procedure. Wound clips were removed 12-14 days post-procedure.

### Analysis of MHC recycling genes

The single-cell RNA sequencing data were obtained from NCBI GEO database GSE168181 [26]. The datasets of naïve LNs and TdLNs were integrated using the “FindIntegrationAnchors” function in the Seurat R package to obtain anchors for batch bias normalization. Subsequently, the “IntegrateData” function was employed to integrate data and correct for technical differences. Next, the top 30 Principal Components (PCs) were selected for Uniform Manifold Approximation and Projection (UMAP) analysis and clustering. Cell types were annotated based on the top highly expressed genes in each cluster. The MHC recycling genes were selected based on a literature search [27–32].

### Biodistribution analysis

2 MDa FITC-dextran (Sigma-Aldrich) and 10 kDa TRITC-dextran (Sigma-Aldrich) in saline were co-injected into locations of interest (forelimb, flank, MFP, hindlimb) in a total volume of 30 µL. Animals were sacrificed 24 hours later, and tissues immediately frozen in optimal cutting temperature (OCT, Tissue-Tek) gel, and then maintained at -80°C. Blocks were then sliced into 10μm sections using a cryostat. Slides were immediately covered with Vectashield (Vector Labs) to avoid diffusion. Vectashield was allowed to harden overnight, and slides were imaged using an Axioscan (Zeiss). Images were processed using ImageJ on a PC using the public domain National Institutes of Health (NIH) Image program (developed at the U.S. NIH and available on the Internet at http://imagej.net/nih-image/).

### Vaccination strategies

Several different vaccination strategies were used in this work: OVA + LPS [10μg OVA (Sigma Aldrich) + 10μg LPS (Sigma-Aldrich) in saline for a total volume of 30µL], LPS alone [10μg LPS (Sigma-Aldrich) in saline for a total volume of 30µL], Titermax [10µL Titermax emulsified with 10µL saline], alum [0.5mg alum (VWR International) in saline for a total volume of 30µL], and lysed cells (MCaP cells from culture, after 5 freeze/thaw cycles via dunking in liquid Nitrogen) combined with 10µg LPS and saline in a total volume of 30µL. Treatments were prepared on the day of vaccination. Treatments were injected intradermally using a 31G insulin needle in the wrist skin of the forelimb contralateral to the tumor location (distant), intradermally in the flank skin ipsilateral to the tumor location (locoregional) or i.t. Pressure was applied on the syringe plunger as the needles were removed from the injection site to avoid any backflow of the solution from the injection site. Saline injected i.t. served as a control.

### Antibody enzyme-linked immunosorbent assay (ELISA)

Serum was collected for antibody ELISAs. In short, blood was taken from animals at sacrifice into 1.5mL uncoated Eppendorf tubes. Blood was allowed to clot at 25°C for 30min and then samples spun down at 1000g for 10min. Serum was removed from the tops of samples and maintained at -80°C. To prepare ELISAs, Nunc MaxiSorp high-affinity plates (Thermo Fisher, Inc.) were incubated with OVA (10μg OVA (Sigma Aldrich) in 100 μL PBS) or PBS alone overnight. Plates were then washed with 300μL PBS + Tween 20 (1% Tween 20 in PBS) three times. Serum (5μL serum in 95μL PBS) samples were then added to plates and allowed to incubate 2 hours at room temperature. Plates were then washed with 100µL PBS three times. Streptavidin conjugated anti-mouse IgG antibodies (1:1000, Southern Biotech) were then added to plates and allowed to incubate for 2 hours at room temperature. Plates were washed with 100µL PBS. Biotinylated horse radish peroxidase (100uL, R&D Biosystems) was then added to each sample and allowed to incubate for approximately 20min at room temperature. Stop solution (50µL, R&D Biosystems) was then added to each sample, and absorbance read at 450nm and 540nm using a multiplex plate reader (MesoScale Discovery).

### Immunofluorescence

Animals were sacrificed, and dissected tissues were placed into OCT gel (Tissue-Tek) and frozen. Tissues were maintained at -80°C. Blocks were then sliced into 10μm slices using a cryostat. Slides were placed in -20°C acetone for 10min, and then air dried. Slides were then washed 3 times for 5 min in PBS + TritonX (PBS with 0.5% TritonX-100). 1% normal donkey serum (Millipore Sigma) in PBS was then added to samples and allowed to incubate for 1 hour at room temperature. Slides were then washed. Primary antibodies (Supplemental Table 2) were added to samples and allowed to incubate overnight at 4°C. Slides were then washed with PBS + TritonX. Secondary antibodies were added along with DAPI counter-stain and allowed to incubate for 1 hour at room temperature in the dark. Slides were washed, and then coverslipped with Vectashield (Vector Labs), and allowed to harden overnight. Slides were then imaged using a Axioscan (Zeiss). Images were analyzed using QuPath-0.4.3 [54].

### Statistical Analysis

Data are represented as the mean accompanied by standard error of the mean (SEM), and statistics (one-way, two-way, or repeated measures (RM) analysis of variance (ANOVA) with Tukey’s post-hoc test for grouped analyses; log-rank test for survival analyses; or t-tests, as indicated in figure legends) were calculated using Prism 10 (GraphPad Software Inc., La Jolla, CA, USA). Statistical significance was defined as p < 0.05, 0.01, 0.005, and 0.001, respectively, unless otherwise specified.

### Study Approvals

All experiments were done in accordance with Massachusetts General Hospital Institutional Animal Care and Use Committee policy, per protocols 2011N000085 and 2004N000123.

## Data Availability

All data are presented within the manuscript, and related data are available upon request.

## Acknowledgments

We thank Peigen Huang, Mark Duquette, Carolyn Smith, Igor Gomes dos Santos, Sonu Subudhi, Taylor Uccello, Anna Khachatryan, Marla Marquez, Julia Linke, and Dennis Jones for their advice, guidance and technical support throughout this study. We thank the NIH Tetramer Core Facility (contract number 75N93020D00005) for providing SIINFEKL tetramers.

## Abbreviations

alum: aluminum hydroxide salt
ANOVA: analysis of variance
APC: antigen-presenting cell
Ax LN: axillary lymph node
cDC1: conventional dendritic cell-1
cDC2: conventional dendritic cell-2
Contra Ax LN: contralateral axillary lymph node
Contra Ing LN: contralateral inguinal lymph node
DC: dendritic cell
dLN: draining lymph node
DMEM: Dulbecco’s modified eagle medium
ELISA: Enzyme-linked immunosorbent assay
FBS: fetal bovine serum
GzmB: granzyme B
HBSS: Hank’s buffered salt solution
ICB: immune checkpoint blockade
IFN: interferon
Ing LN: inguinal lymph node
Ipsi Ax LN: ipsilateral axillary lymph node
Ipsi Ing LN: ipsilateral inguinal lymph node
i.t.: intratumoral
LN: lymph node
LPS: lipopolysaccharide
mAb: monoclonal antibody
MFI: mean fluorescence index
MFP: mammary fatpad
MHC: major histocompatibility complex
NdLN: non-draining lymph node
NIH: National Institutes of Health
n.s.: not significant
OCT: optimal cutting temperature
OVA: ovalbumin
PBS: phosphate-buffered saline
PD1: programmed death-1
pDC: plasmacytoid dendritic cell
RM-ANOVA: repeat measures analysis of variance
RPMI: Roswell Park Memorial Institute
SEM: standard error of the mean
TdLN: tumor-draining lymph node
Titermax: Titermax Gold

## Author contributions

Conceptualization: MJO, TPP, LLM

Formal analysis: MJO

Investigation: MJO, PL, HZ, NCA, LM, JJR, MNR, LL

Writing – Original Draft: MJO

Supervision: TPP, LLM, SC, GMB, JWB, JMU

## Data and materials availability

All data are available in the main text or supplementary materials.

## Supplemental Figure Legends

**Supplemental Figure 1: Gating strategy for surface phenotyping**. Shown in a representative spleen sample.

**Supplemental Figure 2:** (**A**) Frequency of CD11c^+^ cells having processed DQ-OVA after ex vivo stimulation as in Fig. 1B. (**B**) Frequency of CD11c^+^ cells having processed DQ-OVA as a function of OVA surface presentation on MHCI. n.s. = not significant (p>0.05).

**Supplemental Figure 3: Phenotype of locoregional tissue-resident DCs is altered.** CD80^+^ (**A**), MHCI^hi^ (**B**), and MHCII^hi^ (**C**) cells as a frequency of CD11c^+^ cells from tissues of interest from animals with or without day 7 E0771 tumors, after immediate staining upon dissection; MHCI mean fluorescence index (MFI) (**D**) and MHCII MFI (**E**) among all CD11c^+^ cells in relevant tissues in day 7 E0771- bearing animals compared to naïve animals, immediately after dissection from animals. *indicates significance by one-way ANOVA with Tukey’s post-hoc test. * indicates p<0.05, ** indicates p<0.01, **** indicates p<0.001. n = 5 animals.

**Supplemental Figure 4: Phenotype of DCs in day 14, not day 7 (as in Supplemental Figure 3), E0771 tumor-bearing versus naïve animals in LNs and spleens.** CD80^+^ (**A**), MHCI^+^ (**B**), and MHCII^+^ (**C**) cells as a frequency of CD11c^+^ cells in relevant tissues in day 14 E0771-bearing mice, compared to naïve animals. (**D**) CD80^+^, MHCI^+^, and MHCII^+^ cells as a frequency of CD11c^+^ cells in spleens of day 7 or day 14 E0771 tumor-bearing and naïve animals. *indicates significance by one-way ANOVA with Tukey’s post-hoc test. * indicates p<0.05, ** indicates p<0.01, *** indicates p<0.005, **** indicates p<0.001. n = 5 animals.

**Supplemental Figure 5: Expression of genes of interest in MHC recycling in naive LN versus TDLNs based on data generated in** [26]. The dot plot represents the single-cell gene expression of MHC recycling-related genes in 4T1 TdLNs and naïve LNs from immunocompetent Balb/c mice. Gene expression values were scaled and normalized using the Seurat R package. More red colors indicate high expression, while more green colors indicate low expression.

**Supplemental Figure 6: Purity of CD11c^+^ and CD8^+^ cells after isolation, relevant to Fig. 1E-F.** (**A**) Frequency of CD11c^+^ cells of live single cells in the sample after positive magnetic isolation from a representative naïve spleen sample, (**B**) Frequency of CD8^+^ cells of live single cells in the sample after negative magnetic isolation from an OT-I spleen sample. Frequency of OT-I CD8^+^ T cells containing PD1 (**C**) and Tcf1 (**D**) after co-incubation with 100,000 DCs (by CD11c positive magnetic isolation) pre-incubated with OVA for 6 hours from different tissue sources, and then incubated with OT-I cells for 18 hours.

**Supplemental Figure 7: Impaired ability to present neoantigens is dependent on the presence of a tumor, other tissues analyzed.** (**A**) Schematic demonstrating experimental setup and timeline to analyze impacts of tumor resection on antigen processing and presentation. (**B**) Frequency of CD11c^+^ cells presenting SIINFEKL after *ex vivo* stimulation over time after resection of the primary tumor relative to tumor-naïve animals in the Ipsi Ax and Contra Ing LNs (other tissues in Fig. 1G). (**C**) Frequency of CD11c^+^ cells presenting SIINFEKL in different tissues post-primary tumor resection (non-normalized data from Fig. 1G and Supplemental Figure 7B). * indicates significance by two-way ANOVA. p represents deviation from 0 based on simple linear regression (D-E).* indicates p<0.05, ** indicates p<0.01.

**Supplemental Figure 8: CD80 and MHCII levels after tumor resection surgery.** (**A**) Frequency of CD11c^+^ cells expressing CD80 over time after resection of the primary E0771 tumor, relative to tumor-naïve animals, in relevant tissues. Frequency of CD11c^+^ cells expressing CD80 (**B**) and MHCII (**C**) in different tissues post-primary tumor resection (all samples at day 28, including both Met+ and Met-). * indicates significance by two-way ANOVA with Tukey’s post-hoc test. * indicates p<0.05, ** indicates p<0.01, *** indicates p<0.005, **** indicates p<0.001.

**Supplemental Figure 9: Determining IL1**β **is relevant to antigen presentation.** (**A**) Cytokine levels in Ipsi Ing and Contra Ax LNs in either naïve or tumor (E0771, day 7)-bearing animals based on multiplexed ELISA. (**B**) Cells presenting SIINFEKL antigen after ex vivo stimulation with cytokines that were different in Supplemental Figure 9A (same experimental design as Figure 2B). Tcf1 (**C**) and PD1 (**D**) expression after co-culture with DCs from various tissues, as in Fig. 2C. * indicates significance by two-way ANOVA with Tukey’s post-hoc test. * indicates p<0.05, ** indicates p<0.01.

**Supplemental Figure 10: Purity of isolated cells in Fig. 2C.** Purity of CD11c^+^ cells as isolated from a representative spleen by positive magnetic isolation (**A**); and of CD8^+^ cells isolated from an OT-I spleen by negative magnetic isolation (**B**) utilized in experiment in Fig 2C.

**Supplemental Figure 11: Antigen processing and presentation in remaining tissues (relevant to Fig. 2E-L).** Frequency of cells having processed DQ-OVA protein (**A**) or presenting SIINFEKL antigen within the MHCI complex (**B**) after IL1β (1ng at each dosing) delivery within the MFP and spleen of naïve animals. Number of CD11c^+^ cells also having processed DQ-OVA (**C**), CD11c^+^ cells also presenting the SIINFEKL antigen (**D**), and number of CD3^+^CD8^+^ cells specific for the SIINFEKL antigen based on tetramer staining (**E**) in tumor-bearing animals.

**Supplemental Figure 12: Effect of delivery of immunological adjuvant on Ipsi Ax, Contra Ing, Spleen.** (**A**) Frequency of CD11c^+^ cells with SIINFEKL loaded in the MHCI complex, within relevant tissues not displayed in Fig. 3 after delivery of immunological adjuvant. (**B**) Frequency of CD8^+^ T cells tagged with SIINFEKL- specific tetramer in relevant tissues not displayed in Fig. 3 after delivery of immunological adjuvant. * indicates significance by two-way ANOVA with Tukey’s post-hoc test. * indicates p<0.05; ** indicates p<0.01; **** indicates p<0.001.

**Supplemental Figure 13: Anti-OVA IgG in peripheral blood after vaccination.**

Measured in serum collected on day 19 of experimental conditions in Fig 3A. * indicates significance by one-way ANOVA with Tukey’s post-hoc test. ns indicates not significant (p>0.05), * indicates p<0.05; ** indicates p<0.01; **** indicates p<0.001.

**Supplemental Figure 14: Delivering adjuvant to distant LNs slows tumor growth.**

Tumor growth after Titermax (**A**) or alum (**B**) was delivered to E0771-bearing animals according to schematic in Fig. 4A. (**C**) Images representing tumors after Titermax was delivered to E0771-bearing animals according to schematic in Fig. 4A but sacrificed on day 21, stained with CD8 and GzmB and counterstained with DAPI. Number of Tcf7^+^ (**D**), and PD1^+^ (**E**) cells per mm^2^ in tumors after Titermax delivery according to schematic in Fig. 4A with sacrifice on day 21. * indicates significance by repeat measures one-way ANOVA (B) or one-way ANOVA (D,E). * indicates p<0.05. n=6-8 animals per group. Scale bar indicates 1 cm.

**Supplemental Figure 15: Individual growth curves after adjuvant delivery.**

Individual growth curves after Titermax (**A**), or alum (**B**) was delivered to E0771- bearing animals, or alum was delivered to 4T1-bearing animals (**C**) according to timeline described in Fig. 4A but without sacrifice at day 21. n=6-8 animals.

**Supplemental Figure 16: Combining distant delivery of immunological adjuvant with ICB leads to long-term survival and protection from recurrent disease.**

(**A**) Experimental timeline for combination of adjuvant delivery with ICB. Tumor growth (**B**) and survival (**C**) after E0771-OVA tumor-bearing animals were treated with OVA+LPS (or vehicle) in relevant locations and αPD1 mAb (or vehicle) according to experimental design in Supplemental Figure 16A. (**D**) Tumor growth after E0771-OVA cells were implanted in the original tumor location of surviving animals according to the experimental design in Supplemental Figure 16A. (**E**) Tumor growth after parental E0771 cells were implanted into the original MFP of surviving animals according to the experimental design in Supplemental Figure 16A.

**Supplemental Figure 17: Individual growth curves after vaccination combined with ICB.** Individual growth curves after LPS+ OVA was delivered to E0771- bearing animals, followed by ICB (experimental design in Fig 4A but without sacrifice at day 21).

**Supplemental Figure 18:** IL1β concentration in relevant tissues after distant adjuvant delivery (Naive and tumor-bearing data equivalent to Fig. 2A). n=4-8 animals. * indicates significance by two-way ANOVA with Tukey’s post-hoc test. **** indicates p<0.001.

**Supplemental Figure 19: Individual growth curves from IL1**β **studies according to experimental design in Fig. 5A.** n=7-9 animals.

**Supplemental Table 1: Antibodies used in flow cytometry staining. Supplemental Table 2: Antibodies used in immunofluorescence staining.**

