## Supplemental Figures for "Overcoming impaired antigen presentation in tumor draining lymph nodes facilitates immunotherapy"

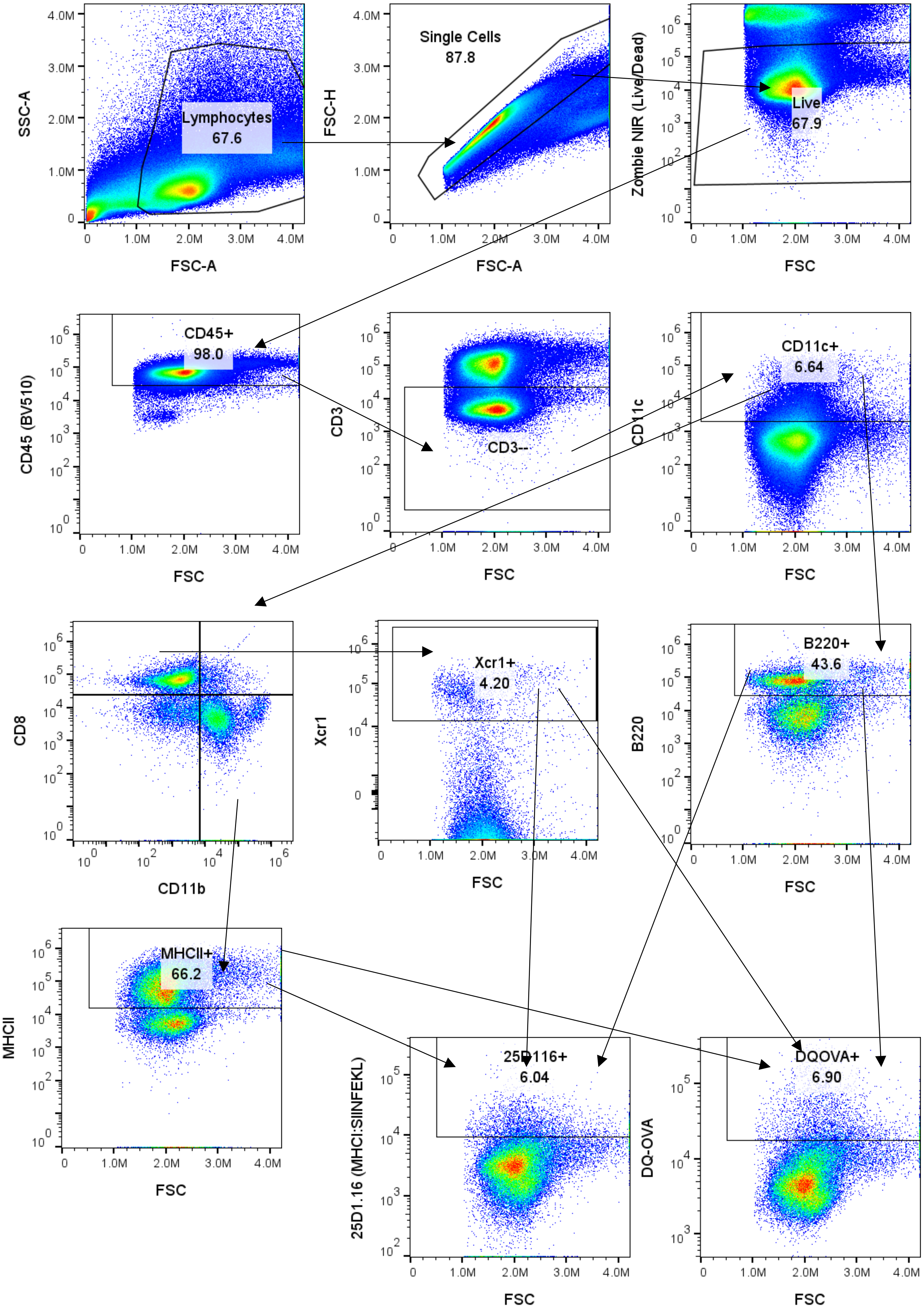

Lymphocytes 67.6
Single Cells 87.8
Live 67.9
CD45+ 98.0
CD3--
CD11c+ 6.64
Xcr1+ 4.20
B220+ 43.6
MHCII+ 66.2
25D116+ 6.04
DQOVA+ 6.90
SSC-A
FSC-A
FSC-H
Zombie NIR (Live/Dead)
FSC
CD45 (BV510)
CD3
CD11c
CD8
CD11b
Xcr1
B220
MHCII
25D1.16 (MHCI:SIINFEKL)
DQ-OVA

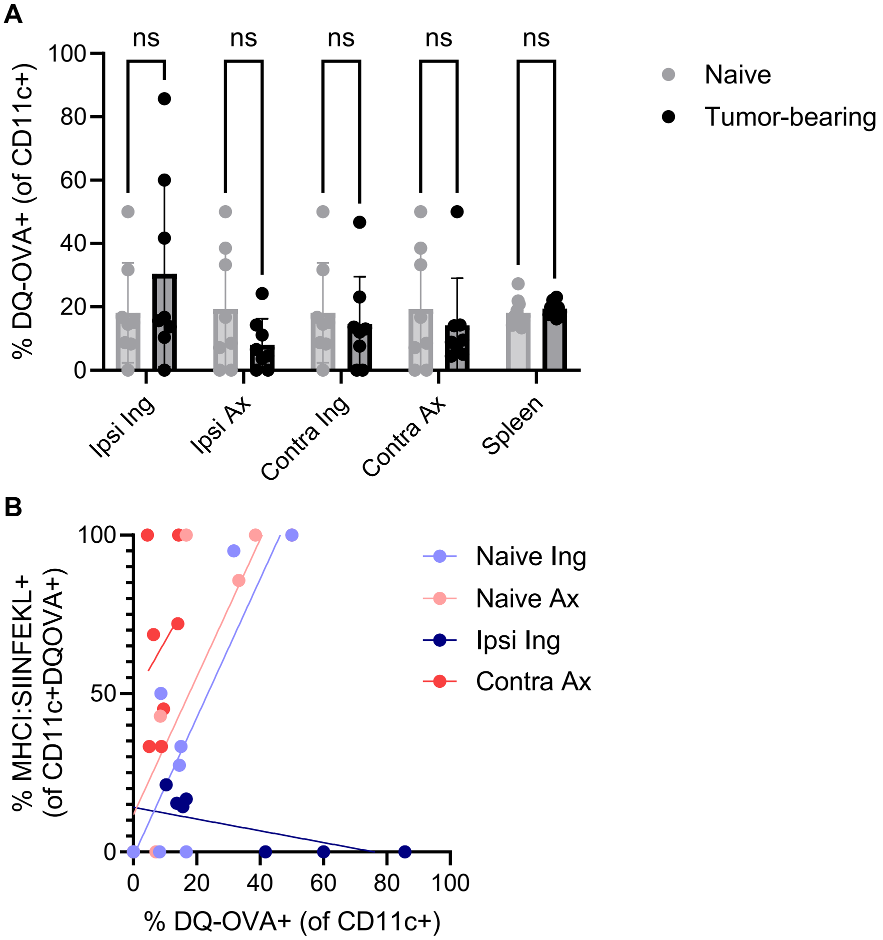
A
ns
% DQ-OVA+ (of CD11c+)
0
20
40
60
80
100
Ipsi Ing
Ipsi Ax
Contra Ing
Contra Ax
Spleen
Naive
Tumor-bearing
B
% MHCI:SIINFEKL+ (of CD11c+DQOVA+)
0
50
100
0
20
40
60
80
100
% DQ-OVA+ (of CD11c+)
Naive Ing
Naive Ax
Ipsi Ing
Contra Ax

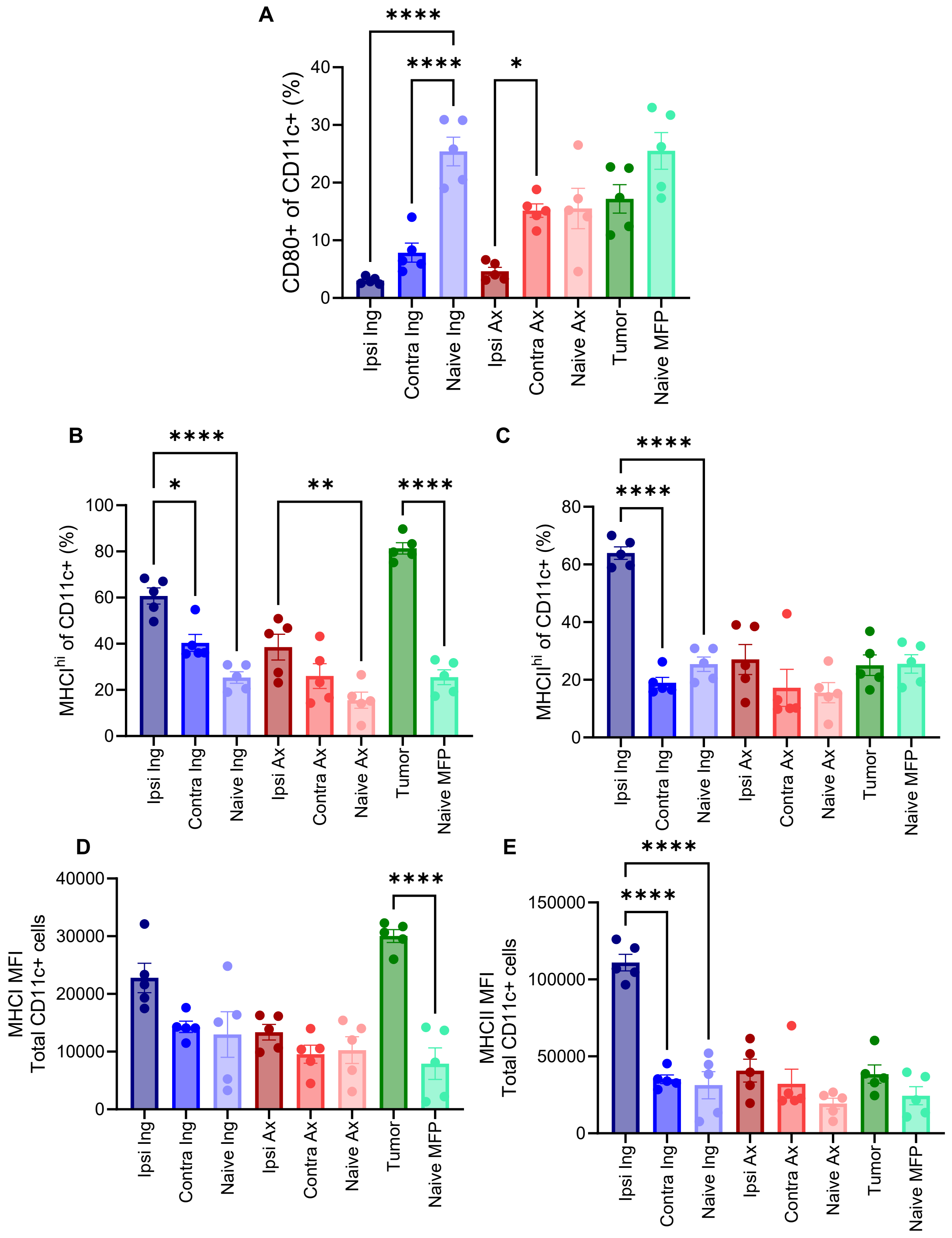
A
B
C
D
E
CD80+ of CD11c+ (%)
MHCI^hi of CD11c+ (%)
MHCII^hi of CD11c+ (%)
MHCI MFI
Total CD11c+ cells
MHCII MFI
Total CD11c+ cells
Ipsi Ing
Contra Ing
Naive Ing
Ipsi Ax
Contra Ax
Naive Ax
Tumor
Naive MFP
****
***
**
*

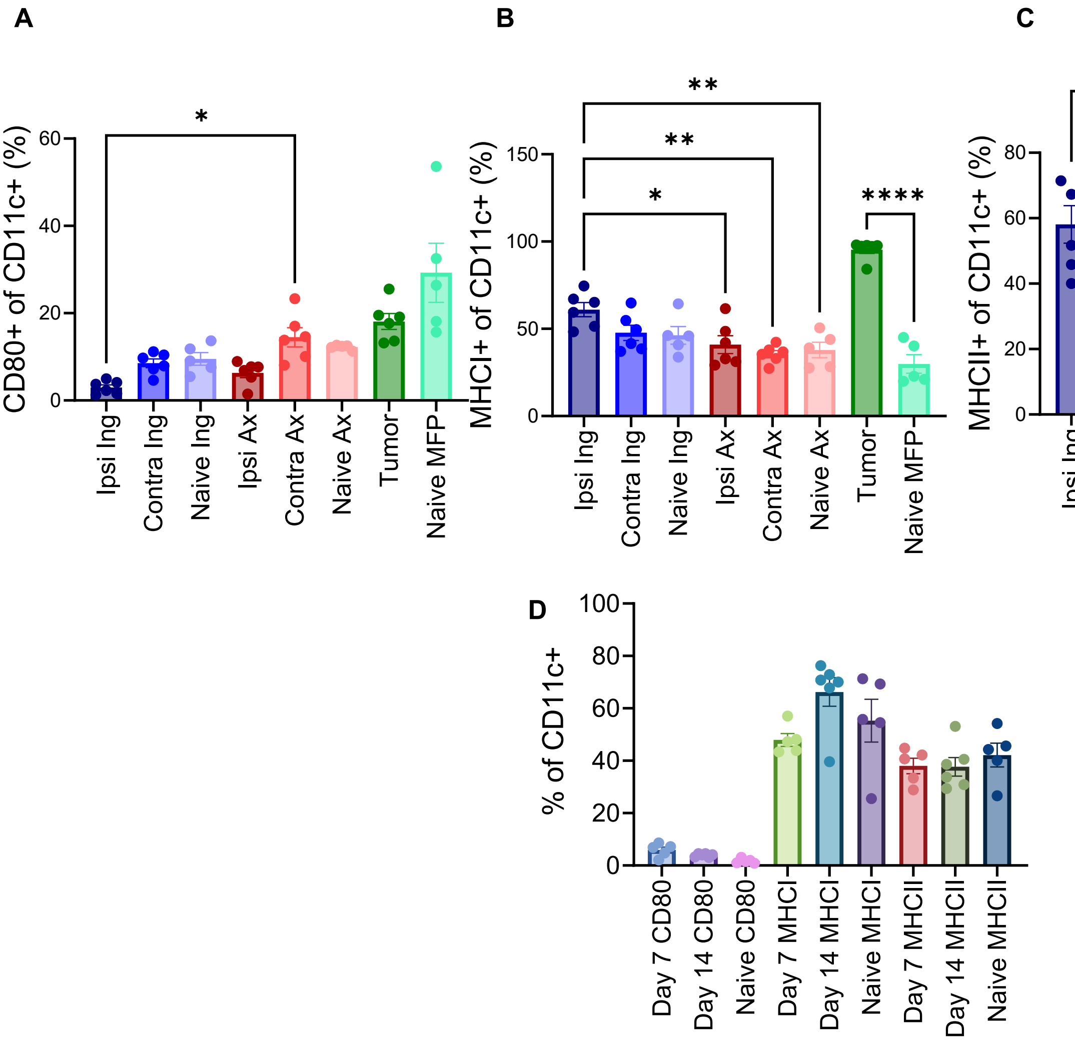
A
CD80+ of CD11c+ (%)
Ipsi Ing
Contra Ing
Naive Ing
Ipsi Ax
Contra Ax
Naive Ax
Tumor
Naive MFP
B
MHCI+ of CD11c+ (%)
C
MHCII+ of CD11c+ (%)
D
% of CD11c+
Day 7 CD80
Day 14 CD80
Naive CD80
Day 7 MHCI
Day 14 MHCI
Naive MHCI
Day 7 MHCII
Day 14 MHCII
Naive MHCII

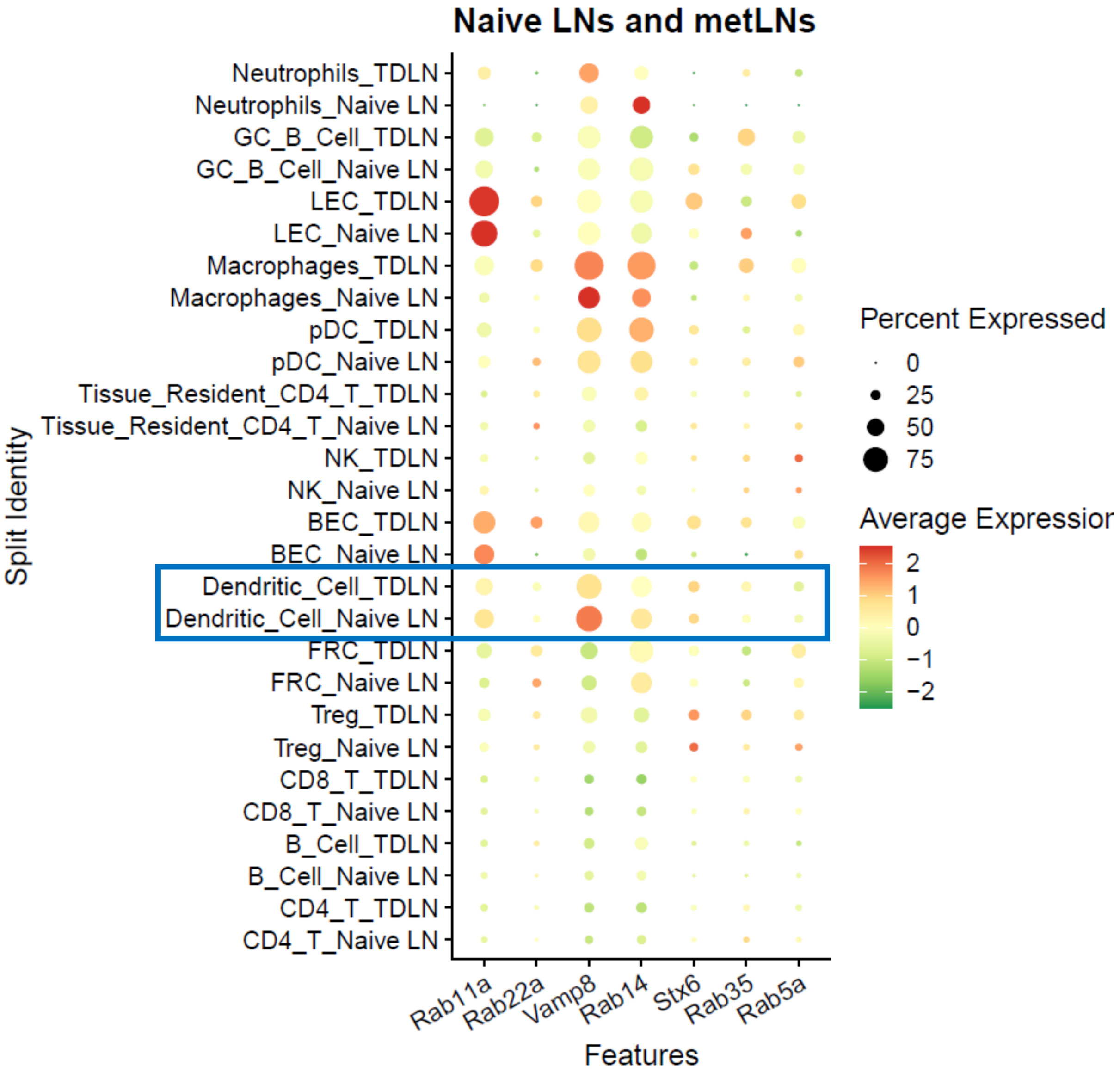

Naive LNs and metLNs
Neutrophils_TDLN
Neutrophils_Naive LN
GC_B_Cell_TDLN
GC_B_Cell_Naive LN
LEC_TDLN
LEC_Naive LN
Macrophages_TDLN
Macrophages_Naive LN
pDC_TDLN
pDC_Naive LN
Tissue_Resident_CD4_T_TDLN
Tissue_Resident_CD4_T_Naive LN
NK_TDLN
NK_Naive LN
BEC_TDLN
BEC_Naive LN
Dendritic_Cell_TDLN
Dendritic_Cell_Naive LN
FRC_TDLN
FRC_Naive LN
Treg_TDLN
Treg_Naive LN
CD8_T_TDLN
CD8_T_Naive LN
B_Cell_TDLN
B_Cell_Naive LN
CD4_T_TDLN
CD4_T_Naive LN
Split Identity
Rab11a
Rab22a
Vamp8
Rab14
Stx6
Rab35
Rab5a
Features
Percent Expressed
0
25
50
75
Average Expressior
2
1
0
−1
−2

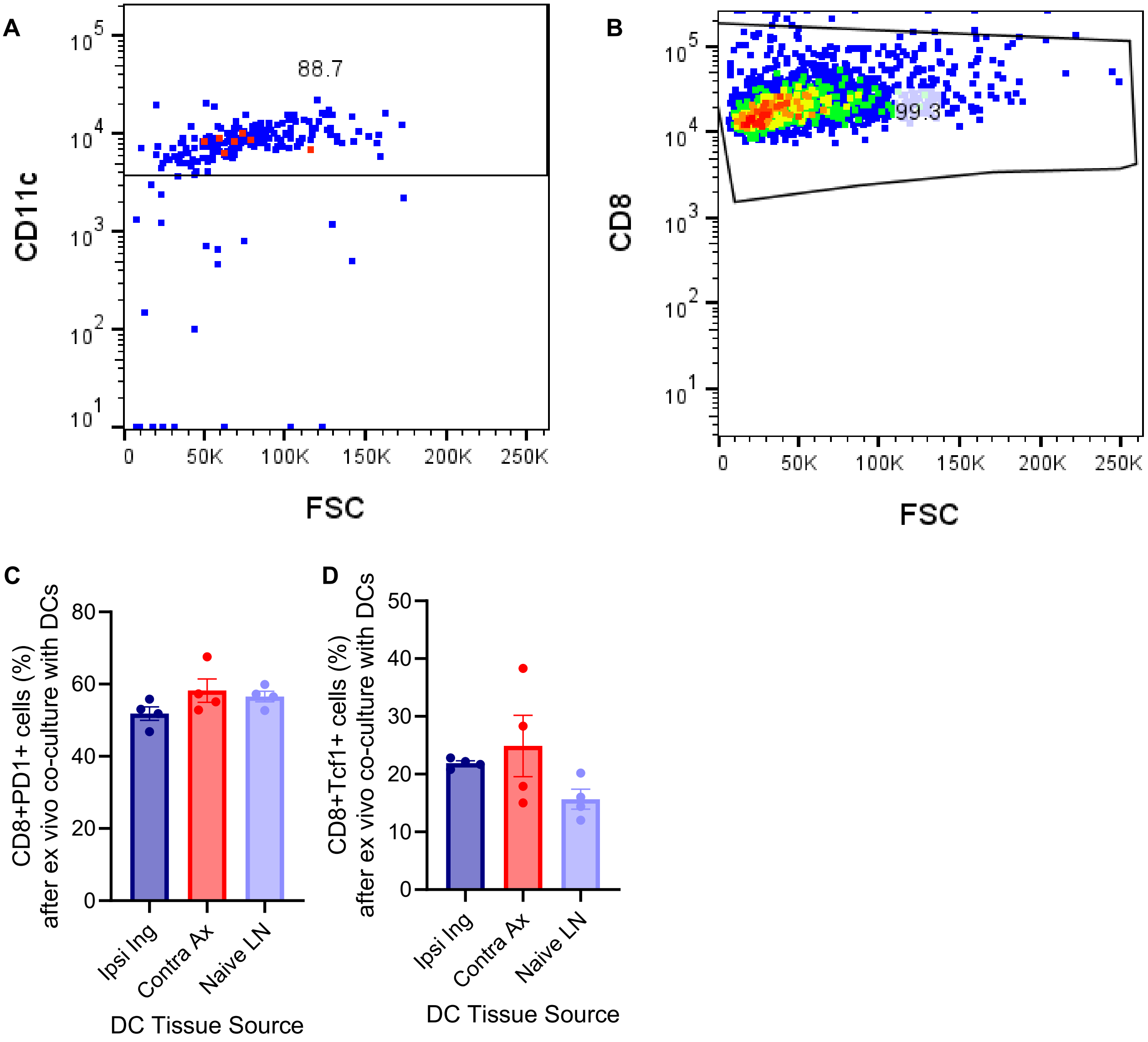

A
88.7
CD11c
FSC
B
99.3
CD8
FSC
C
CD8+PD1+ cells (%) after ex vivo co-culture with DCs
Ipsi Ing
Contra Ax
Naive LN
DC Tissue Source
D
CD8+Tcf1+ cells (%) after ex vivo co-culture with DCs
Ipsi Ing
Contra Ax
Naive LN
DC Tissue Source

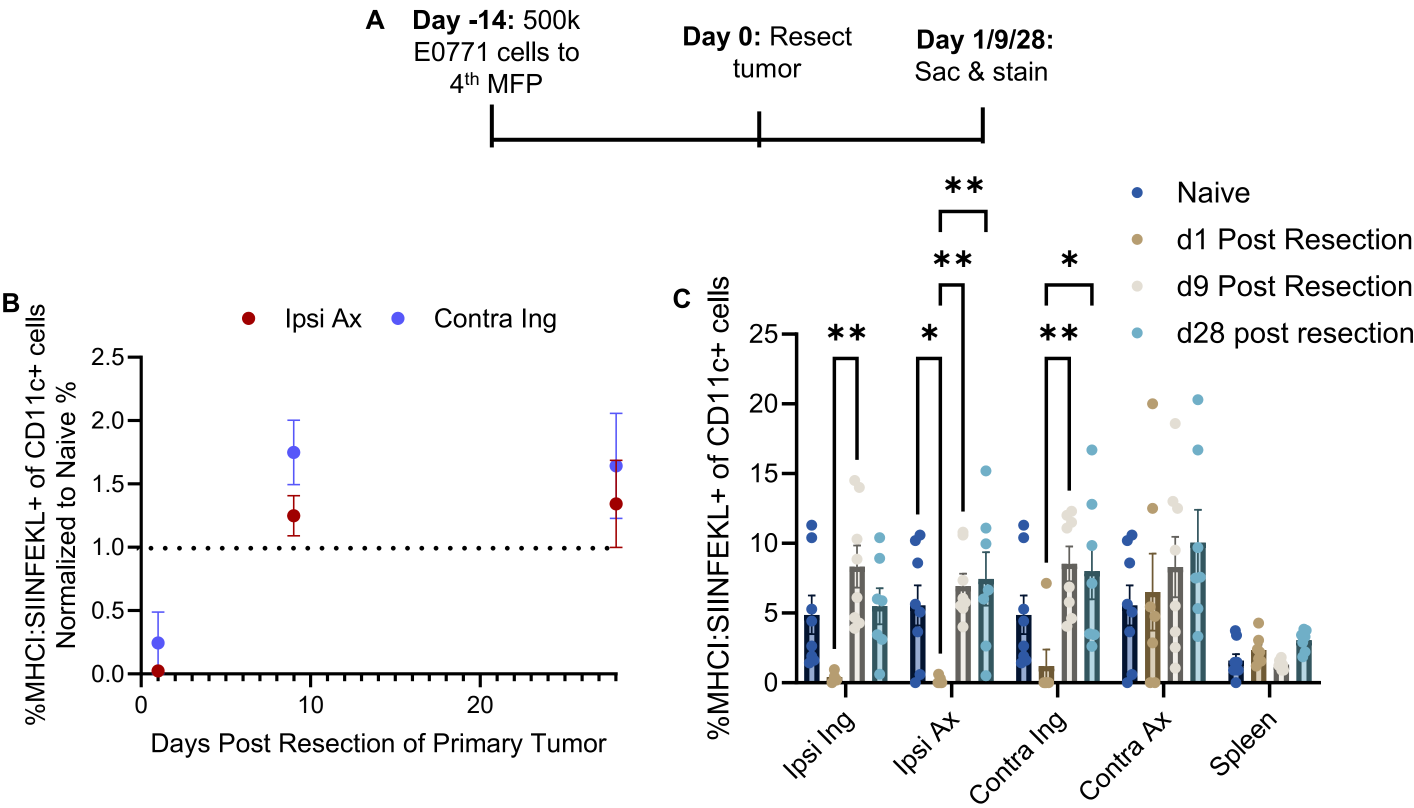
A
Day -14: 500k E0771 cells to 4th MFP
Day 0: Resect tumor
Day 1/9/28: Sac & stain
B
Ipsi Ax
Contra Ing
%MHCI:SIINFEKL+ of CD11c+ cells Normalized to Naive %
Days Post Resection of Primary Tumor
C
Naive
d1 Post Resection
d9 Post Resection
d28 post resection
%MHCI:SIINFEKL+ of CD11c+ cells
Ipsi Ing
Ipsi Ax
Contra Ing
Contra Ax
Spleen

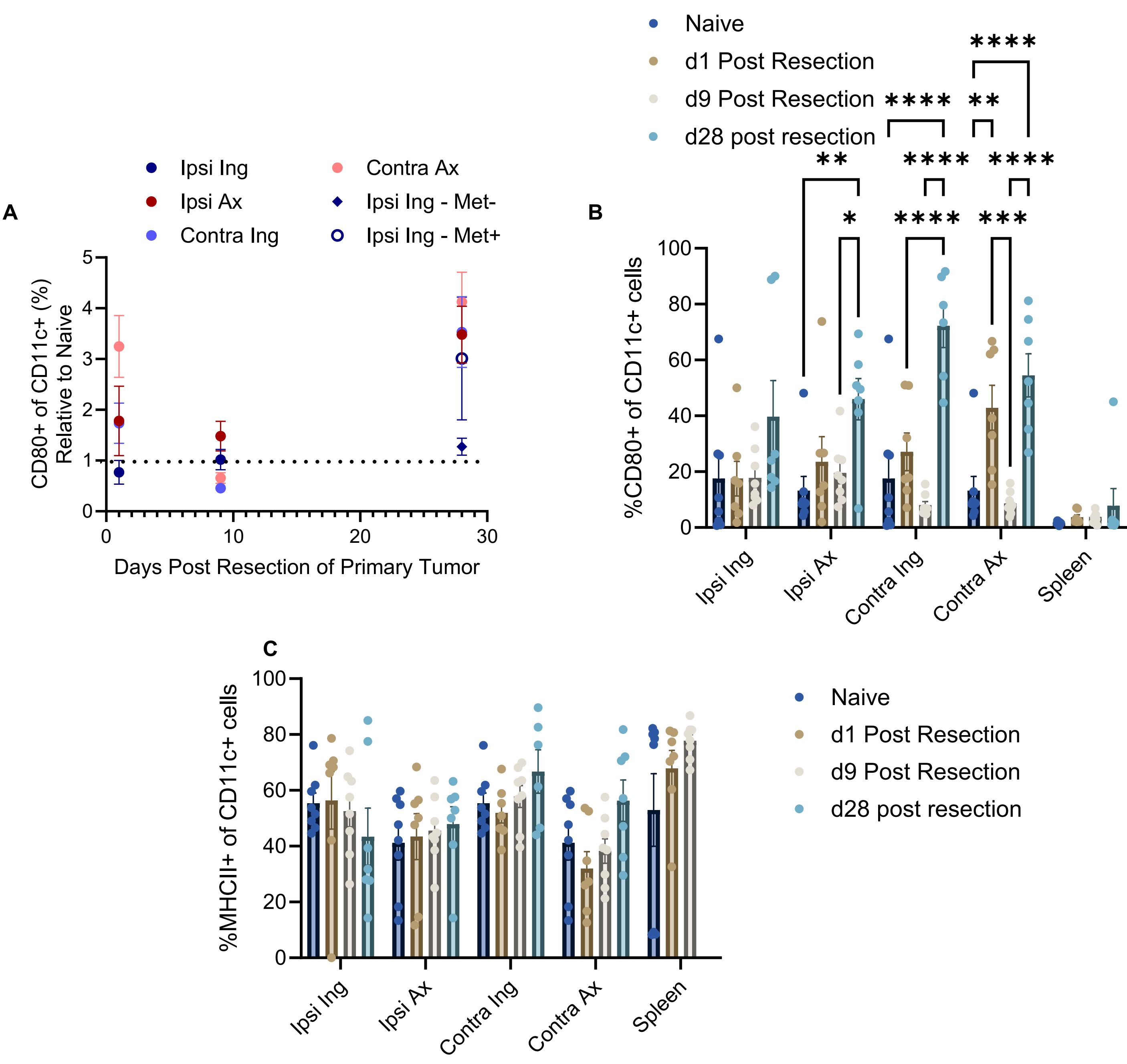

A
Ipsi Ing
Ipsi Ax
Contra Ing
Contra Ax
Ipsi Ing - Met-
Ipsi Ing - Met+
CD80+ of CD11c+ (%) Relative to Naive
Days Post Resection of Primary Tumor
B
Naive
d1 Post Resection
d9 Post Resection
d28 post resection
%CD80+ of CD11c+ cells
****
**
*
***
Ipsi Ing
Ipsi Ax
Contra Ing
Contra Ax
Spleen
C
%MHCII+ of CD11c+ cells
Naive
d1 Post Resection
d9 Post Resection
d28 post resection
Ipsi Ing
Ipsi Ax
Contra Ing
Contra Ax
Spleen

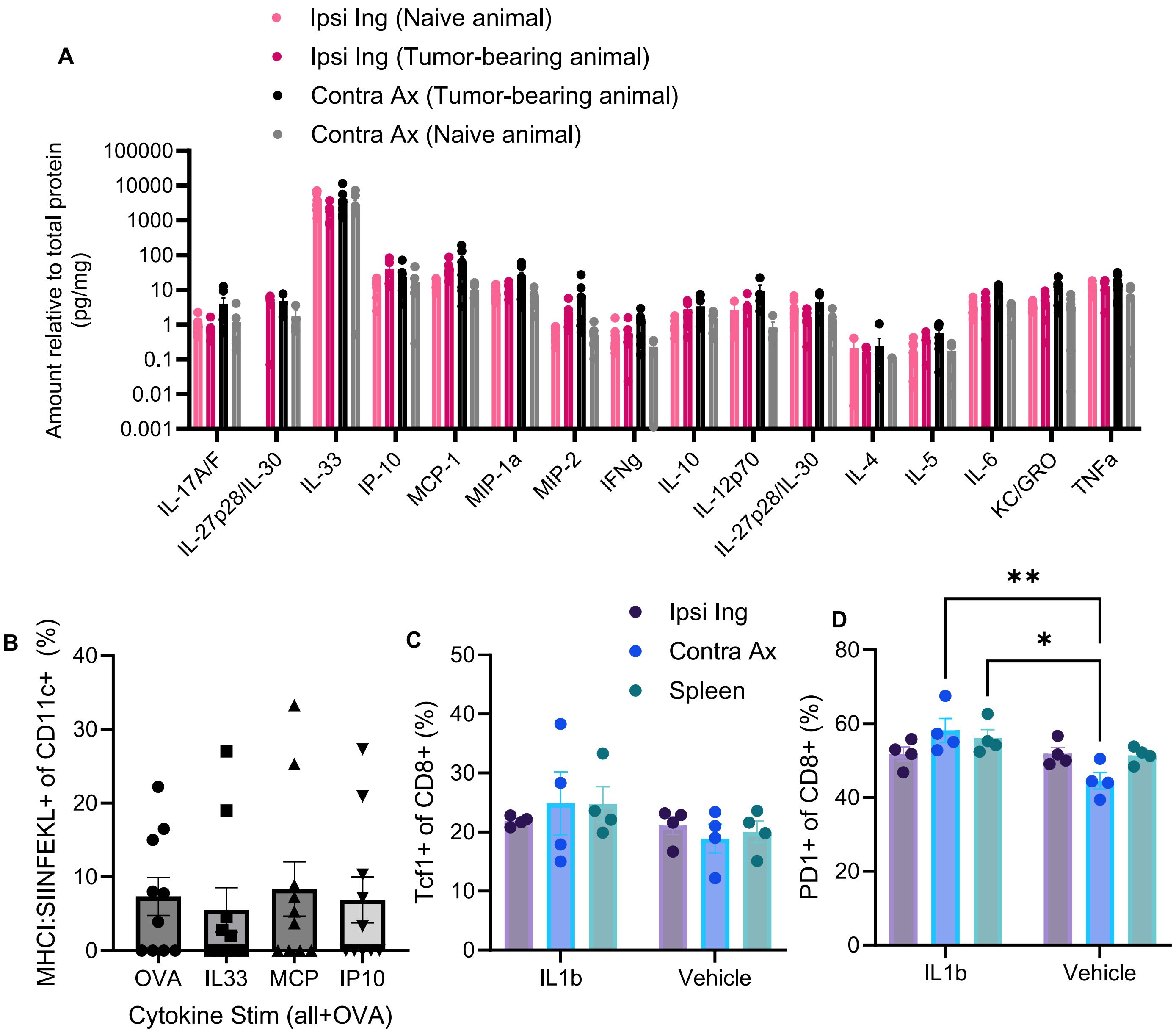

A
Ipsi Ing (Naive animal)
Ipsi Ing (Tumor-bearing animal)
Contra Ax (Tumor-bearing animal)
Contra Ax (Naive animal)
Amount relative to total protein (pg/mg)
100000
10000
1000
100
10
1
0.1
0.01
0.001
IL-17A/F
IL-27p28/IL-30
IL-33
IP-10
MCP-1
MIP-1a
MIP-2
IFNg
IL-10
IL-12p70
IL-27p28/IL-30
IL-4
IL-5
IL-6
KC/GRO
TNFa
B
MHCI:SIINFEKL+ of CD11c+ (%)
40
30
20
10
0
OVA
IL33
MCP
IP10
Cytokine Stim (all+OVA)
C
Tcf1+ of CD8+ (%)
50
40
30
20
10
0
IL1b
Vehicle
Ipsi Ing
Contra Ax
Spleen
D
PD1+ of CD8+ (%)
80
60
40
20
0
**
*
IL1b
Vehicle

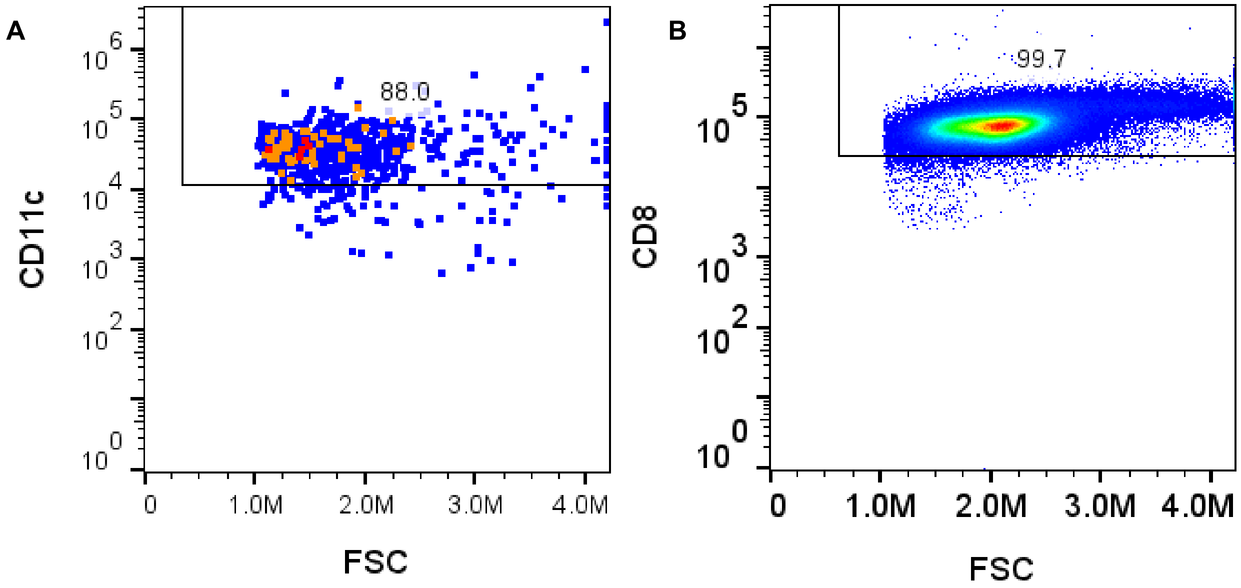
A
88.0
CD11c
FSC
0
1.0M
2.0M
3.0M
4.0M
B
99.7
CD8
FSC
0
1.0M
2.0M
3.0M
4.0M

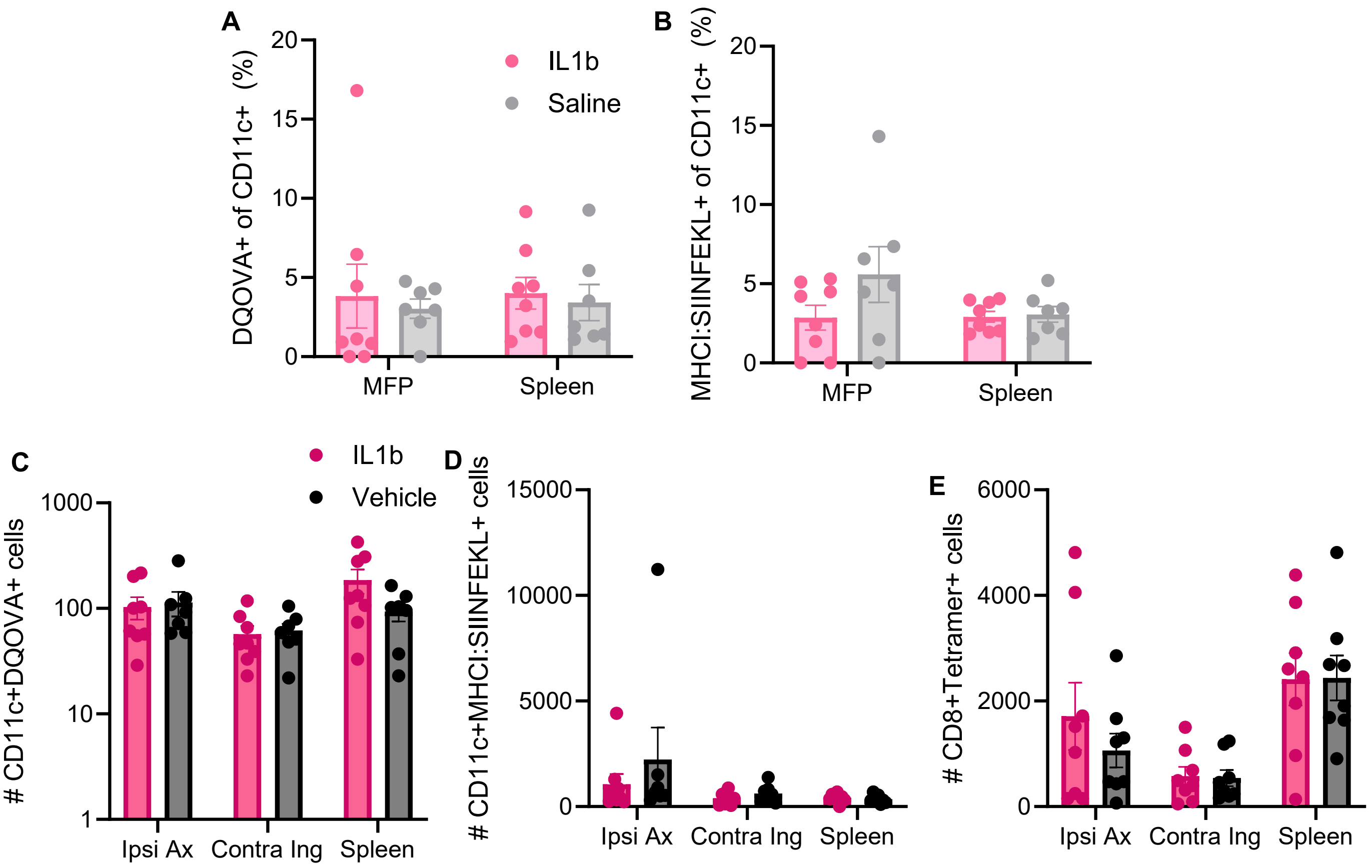
A
B
C
D
E
IL1b
Saline
Vehicle
DQOVA+ of CD11c+ (%)
MHCI:SIINFEKL+ of CD11c+ (%)
### CD11c+DQOVA+ cells
### CD11c+MHCI:SIINFEKL+ cells
### CD8+Tetramer+ cells
MFP
Spleen
Ipsi Ax
Contra Ing

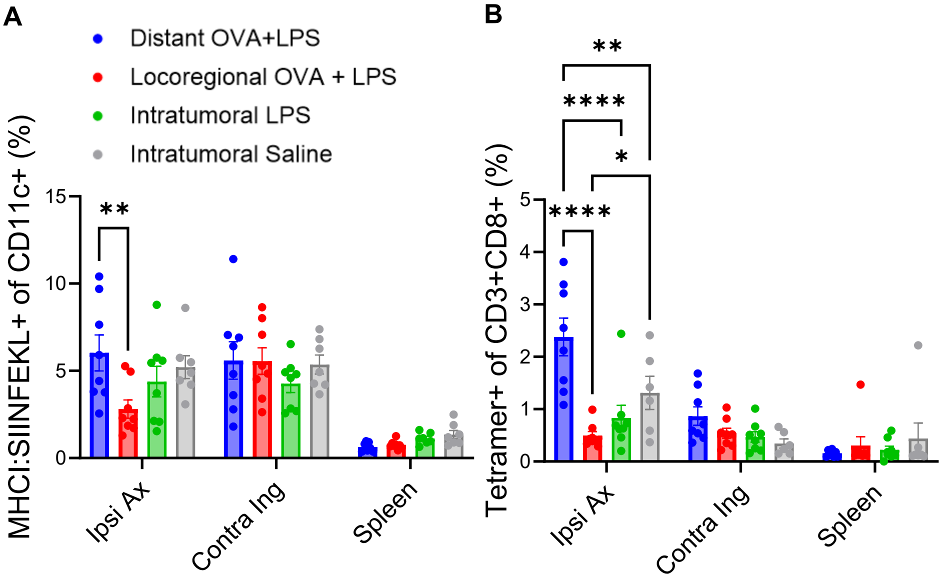
A
B
Distant OVA+LPS
Locoregional OVA + LPS
Intratumoral LPS
Intratumoral Saline
MHCI:SIINFEKL+ of CD11c+ (%)
Tetramer+ of CD3+CD8+ (%)
Ipsi Ax
Contra Ing
Spleen
**
****
*

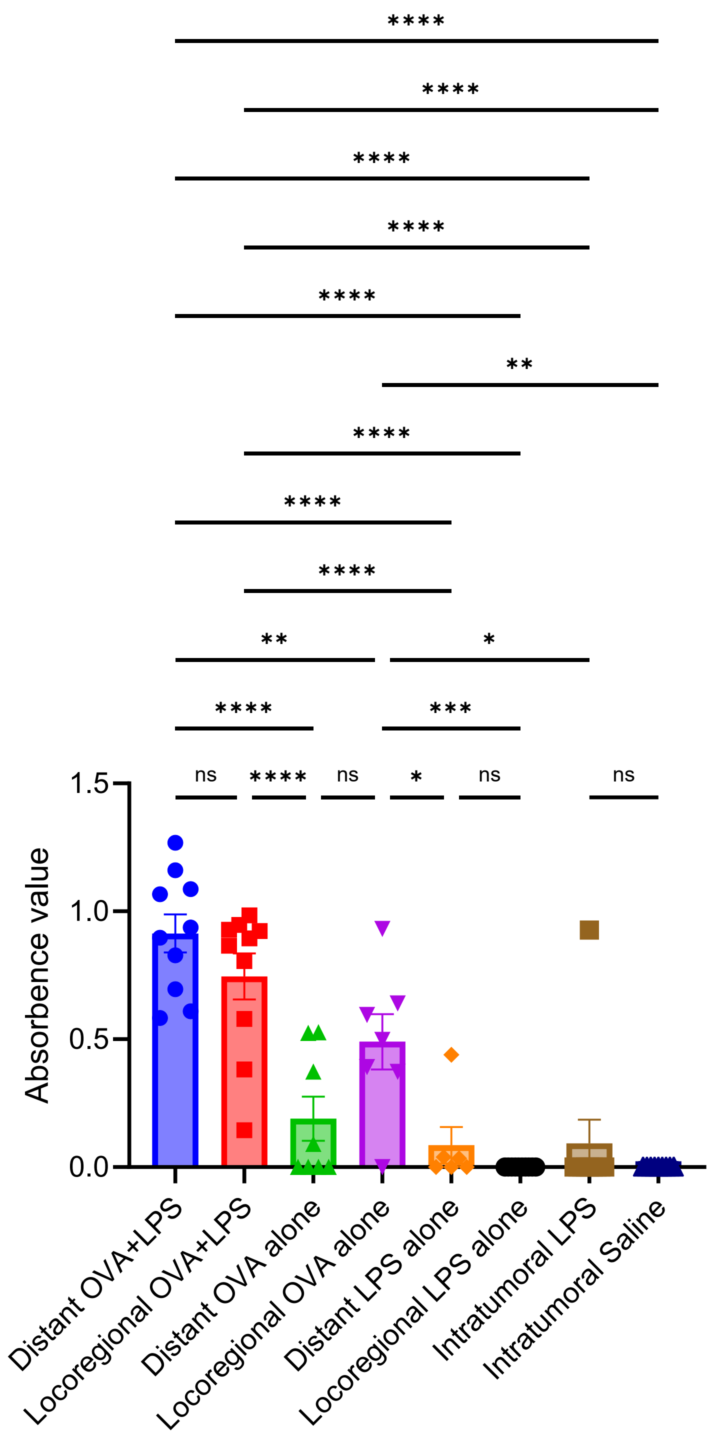

****
****
****
****
****
**
****
****
****
**
*
****
***
ns
****
ns
*
ns
ns
1.5
1.0
0.5
0.0
Absorbence value
Distant OVA+LPS
Locoregional OVA+LPS
Distant OVA alone
Locoregional OVA alone
Distant LPS alone
Locoregional LPS alone
Intratumoral LPS
Intratumoral Saline

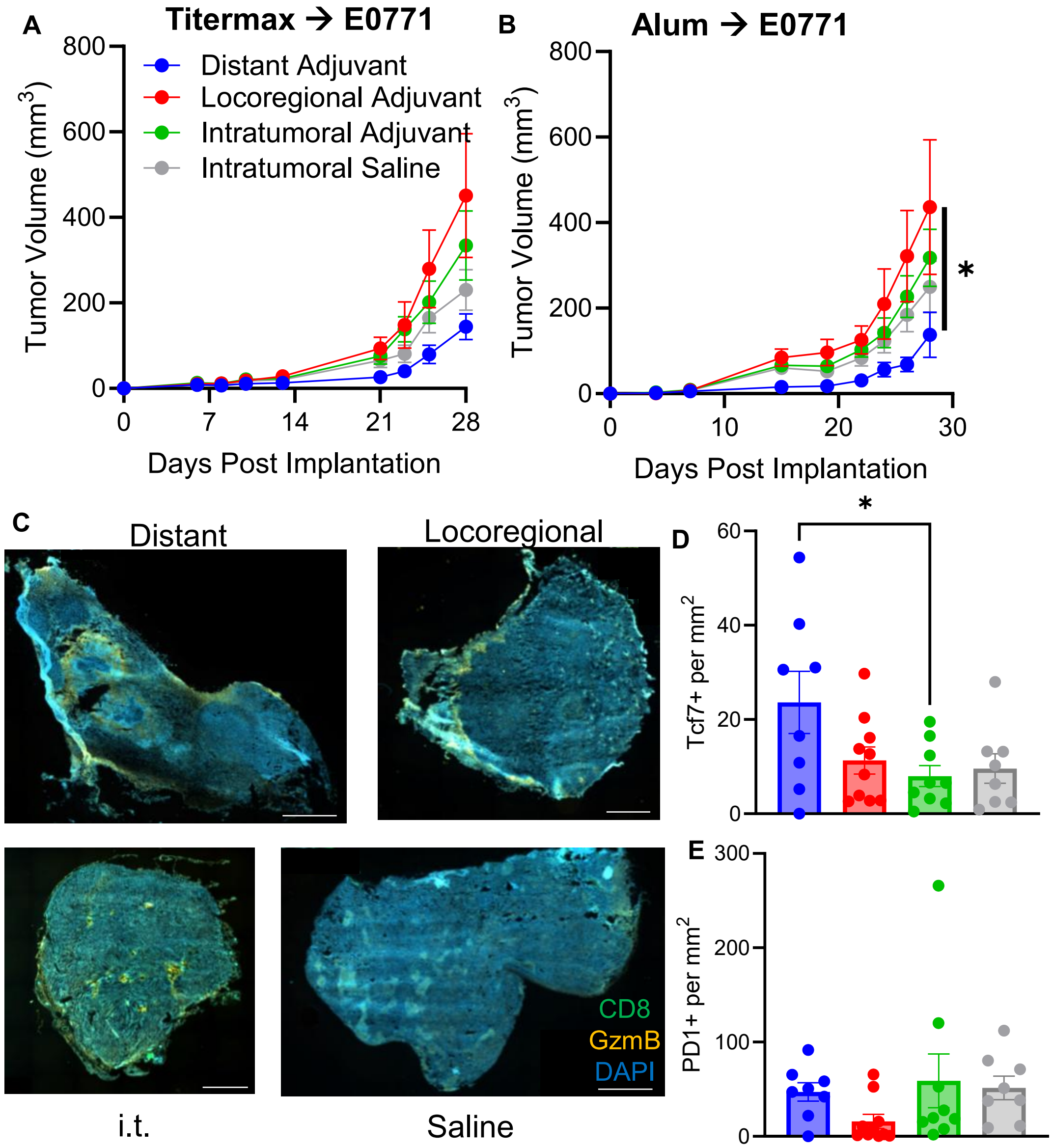
A
Titermax → E0771
Tumor Volume (mm3)
Distant Adjuvant
Locoregional Adjuvant
Intratumoral Adjuvant
Intratumoral Saline
Days Post Implantation
B
Alum → E0771
Tumor Volume (mm3)
*
Days Post Implantation
C
Distant
Locoregional
i.t.
Saline
CD8
GzmB
DAPI
D
Tcf7+ per mm2
*
E
PD1+ per mm2

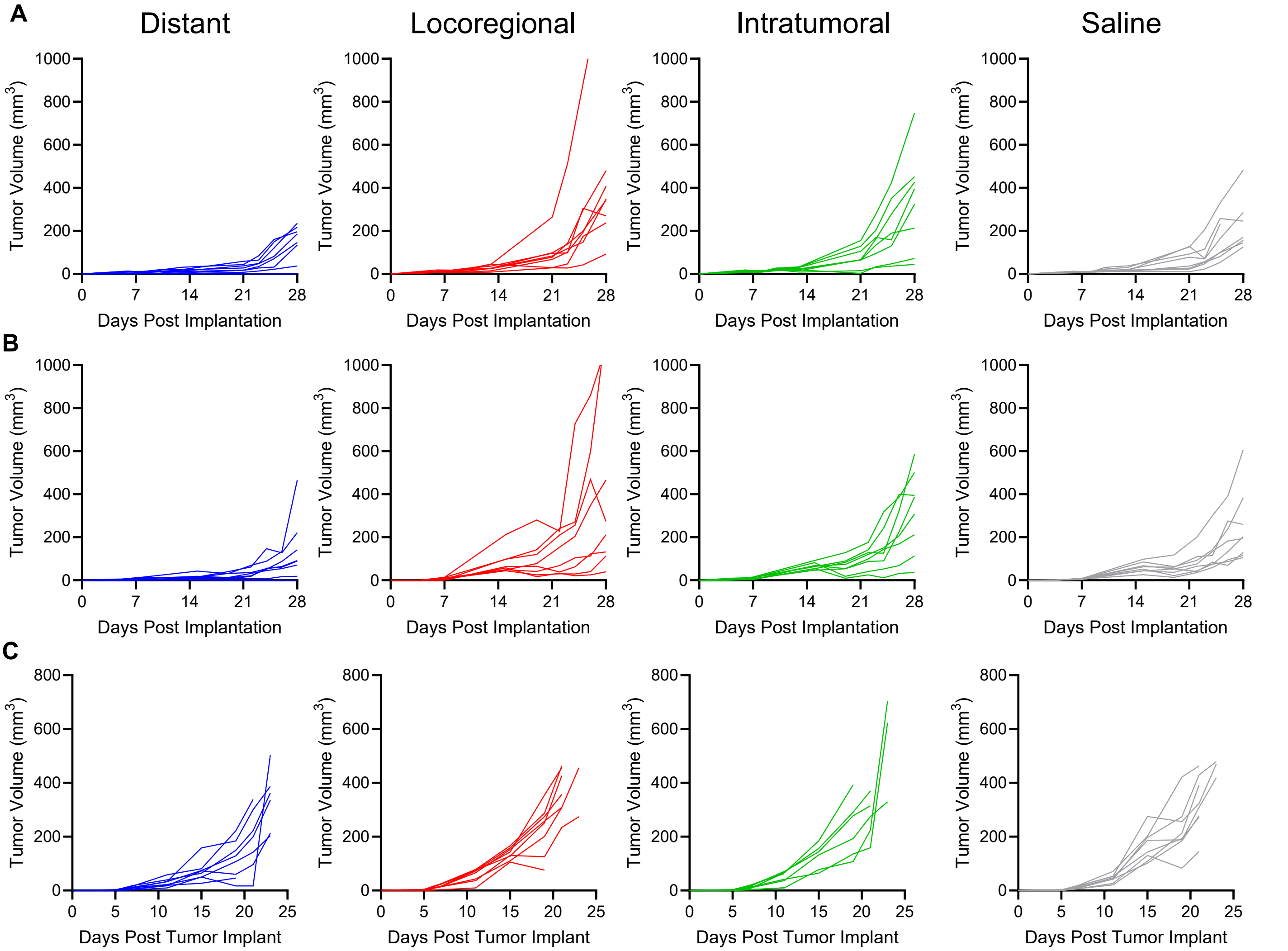

A
Distant
Locoregional
Intratumoral
Saline
Tumor Volume (mm3)
Days Post Implantation
B
C
Days Post Tumor Implant

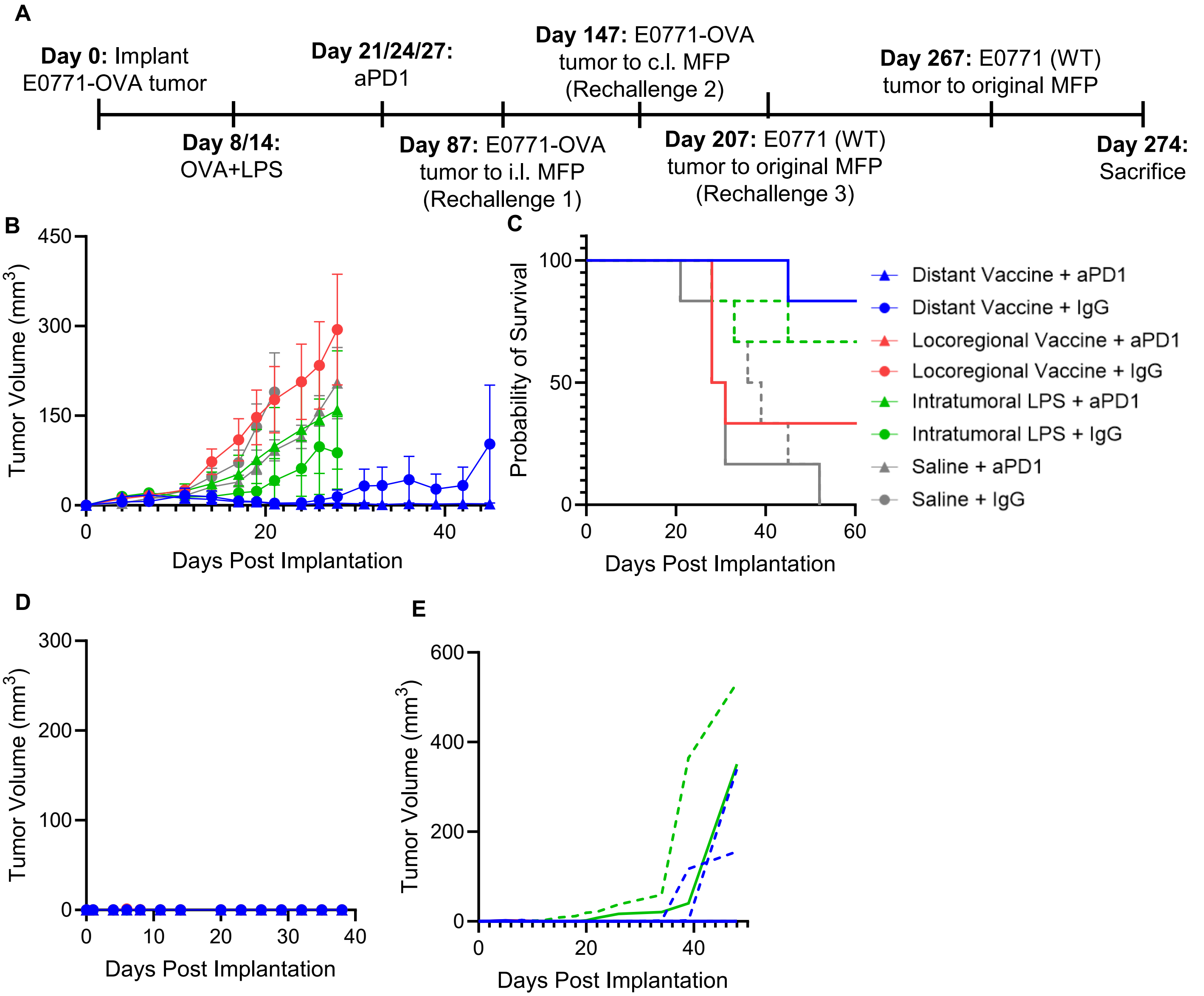
A
Day 0: Implant E0771-OVA tumor
Day 8/14: OVA+LPS
Day 21/24/27: aPD1
Day 87: E0771-OVA tumor to i.l. MFP (Rechallenge 1)
Day 147: E0771-OVA tumor to c.l. MFP (Rechallenge 2)
Day 207: E0771 (WT) tumor to original MFP (Rechallenge 3)
Day 267: E0771 (WT) tumor to original MFP
Day 274: Sacrifice
B
Tumor Volume (mm3)
Days Post Implantation
C
Probability of Survival
Days Post Implantation
Distant Vaccine + aPD1
Distant Vaccine + IgG
Locoregional Vaccine + aPD1
Locoregional Vaccine + IgG
Intratumoral LPS + aPD1
Intratumoral LPS + IgG
Saline + aPD1
Saline + IgG
D
Tumor Volume (mm3)
Days Post Implantation
E
Tumor Volume (mm3)
Days Post Implantation

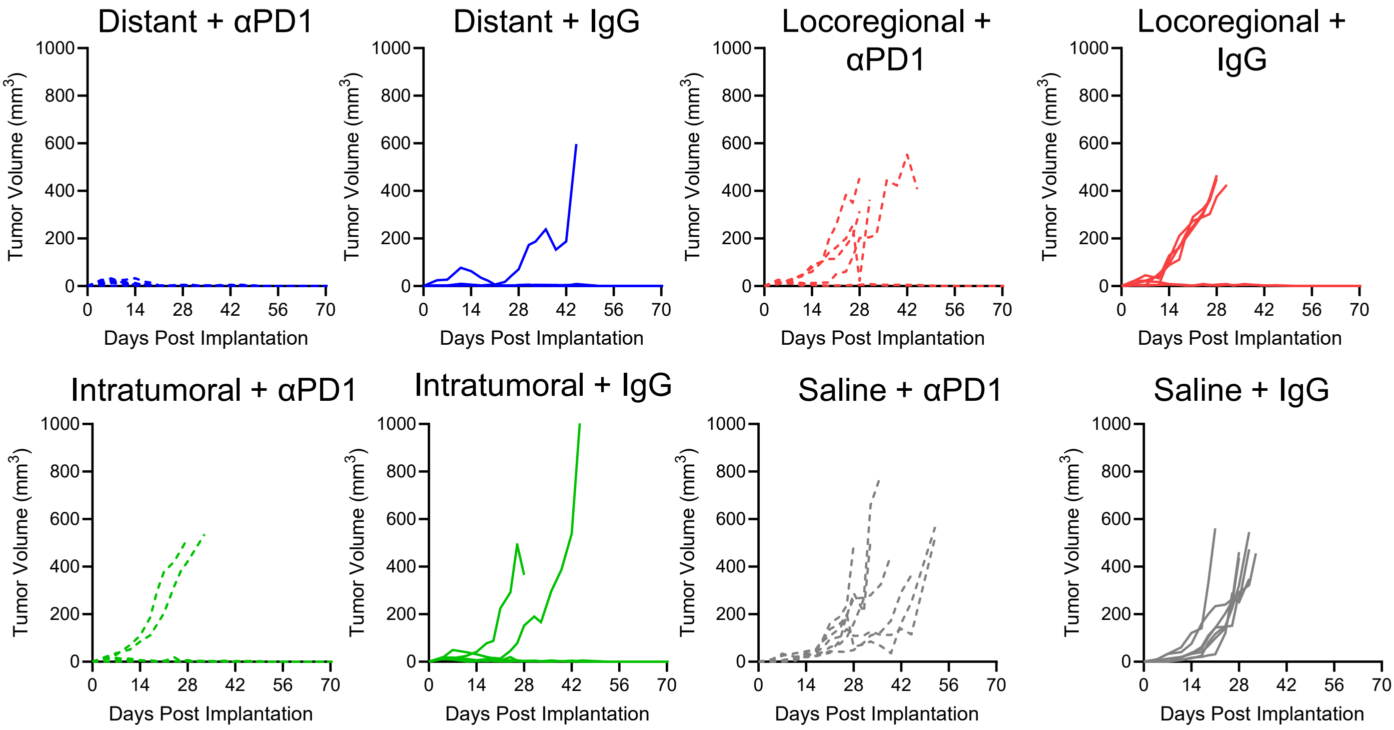

Distant + αPD1
Distant + IgG
Locoregional + αPD1
Locoregional + IgG
Intratumoral + αPD1
Intratumoral + IgG
Saline + αPD1
Saline + IgG
Tumor Volume (mm3)
Days Post Implantation
0
200
400
600
800
1000
0
14
28
42
56
70

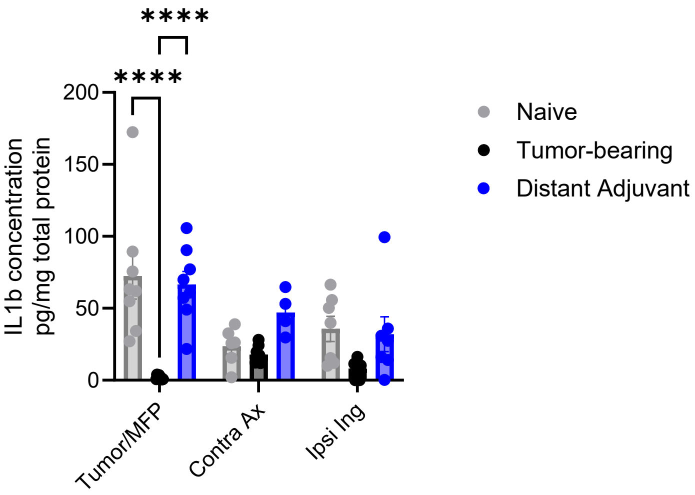

****
****
200
150
100
50
0
IL1b concentration pg/mg total protein
Tumor/MFP
Contra Ax
Ipsi Ing
Naive
Tumor-bearing
Distant Adjuvant

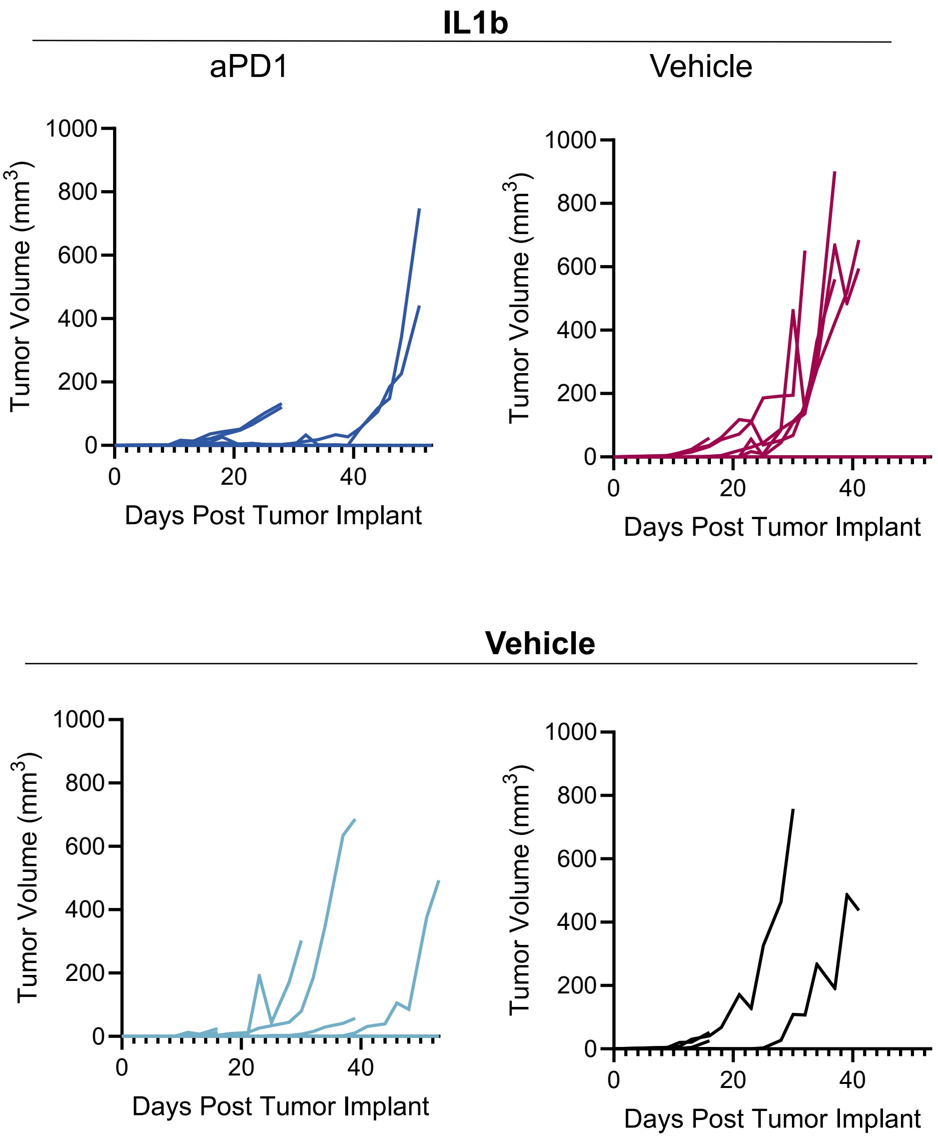

IL1b
aPD1
Vehicle
Vehicle
Tumor Volume (mm3)
Days Post Tumor Implant
1000
800
600
400
200
0
0
20
40

| Marker | Color | Clone | Manufacturer | Cat. No. |
|---|---|---|---|---|
| | | DQ-OVA panel | | |
| CD45 | PerCP | 30-F11 | Biolegend | 103130 |
| CD11c | APC | N418 | Biolegend | 117310 |
| MHCI:SIINFEKL | PE | 25D1.16 | Biolegend | 141604 |
| CD86 | Alexa Fluor 700 | GL-1 | Biolegend | 105024 |
| CD80 | PE-Cy7 | 16-10A1 | Biolegend | 104734 |
| CD206 | APC-Cy7 | 15-2 | Biolegend | 321120 |
| Viability | Zombie Red | | Biolegend | 423110 |
| | | Persistence panel | | |
| CD45 | PerCP | 30-F11 | Biolegend | 103130 |
| CD11c | APC-Cy7 | N418 | Biolegend | 117324 |
| MHCI | Alexa Fluor 647 | 34-1-2S | Biolegend | 114712 |
| MHCII | PE | M5/114.15.2 | Biolegend | 107608 |
| CD86 | Alexa Fluor 700 | GL-1 | Biolegend | 105024 |
| CD80 | PE-Cy7 | 16-10A1 | Biolegend | 104734 |
| CD8 | FITC | 53-6.7 | Biolegend | 100706 |
| Viability | Zombie Red | | Biolegend | 423110 |
| | | IL1β T cell panel | | |
| CD3 | PerCP | 145-2C11 | Biolegend | 100326 |
| CD8 | FITC | 53-6.7 | Biolegend | 100706 |
| PD1 | APC | 29F.1A12 | Biolegend | 135210 |
| CD44 | PE-Cy7 | IM7 | Biolegend | 103030 |
| Tcf1 | PE | 7F11A10 | Biolegend | 655208 |
| GzmB | Alexa Fluor 700 | QA16A02 | Biolegend | 372222 |
| Viability | Zombie Red | | Biolegend | 423110 |
| | | Vaccine T cell panel | | |
| CD3 | PerCP | 145-2C11 | Biolegend | 100326 |
| CD8 | FITC | 53-6.7 | Biolegend | 100706 |
| GzmB | Pacific Blue | GB11 | Biolegend | 515408 |
| PD1 | APC-Cy7 | 29F.1A12 | Biolegend | 135224 |
| Tcf1 | PE | 7F11A10 | Biolegend | 655208 |
| Tim3 | PE-Cy7 | B8.2C12 | Biolegend | 134010 |
| Tetramer | APC | anti-SIINFEKL | NIH Tetramer Core | |
| Viability | Zombie UV | | Biolegend | 423108 |
| | | Vaccine APC panel | | |
| CD45 | APC-Cy7 | QA17A26 | Biolegend | 157618 |
| CD11c | FITC | N418 | Biolegend | 117306 |
| CD80 | PE-Cy7 | 16-10A1 | Biolegend | 104734 |
| MHCI:SIINFEKL | PE | 25D1.16 | Biolegend | 141604 |
| CD19 | APC | 6D5 | Biolegend | 115512 |
| Viability | Zombie UV | | Biolegend | 423108 |
| | | Expanded Ag processing and presentation | | |
| CD45 | BV510 | S18009F | Biolegend | 157219 |
| CD3 | AF488 | 17A2 | Biolegend | 100210 |
| B220 | AF594 | RA3-6B2 | Biolegend | 103254 |
| CD11c | APC | N418 | Biolegend | 117310 |
| MHCI:SIINFEKL | PE | 25D1.16 | Biolegend | 141604 |
| CD11b | BV711 | M1/70 | Biolegend | 101242 |
| F4/80 | PerCP-Cy5.5 | BM8 | Biolegend | 123128 |
| Xcr1 | APC-Cy7 | ZET | Biolegend | 148224 |
| MHCII (I-A/I-E) | AF700 | M5/114.15.2 | Biolegend | 107622 |
| Siglec-H | Pac Blue | 551 | Biolegend | 129610 |
| CD8 | BV421 | 53-6.7 | Biolegend | 100738 |
| CD103 | PE-Cy7 | 2E7 | Biolegend | 121426 |
| Viability | NIR | | Biolegend | 423106 |

| Stain series | Primary target | Primary mAb category # | Secondary target | Secondary Ab color |
|---|---|---|---|---|
| CD8/GzmB | CD8 | abcam ab22378 | Rat | AF488 |
| | GzmB | abcam ab4059 | Rabbit | Cy3 |
| CD8/Tcf7/PD1 | CD8 | abcam ab22378 | Rat | AF488 |
| | Tcf7 | Cell Signalling 2203S | Rabbit | Cy3 |
| | PD1 | eBioscience 14-9985085 | Armenian hamster | AF647 |
